# Sleep deprivation impairs information processing via dysregulation of chloride homeostasis in the prefrontal cortex

**DOI:** 10.64898/2026.03.16.712106

**Authors:** Roberto Frau, Luca Concas, Giulia Braccagni, Francesco Traccis, Caterina Branca, Sara Salviati, Valeria Serra, Eleonora Corridori, Gabriele Nardi, Giacomo Pasquini, Silvia Landi, Michele Santoni, Paolo Follesa, Federico Brandalise, Monica Puligheddu, Miriam Melis, Gian Michele Ratto, Simona Scheggi, Marco Bortolato

## Abstract

Sleep deprivation (SD) impairs information processing through alterations of prefrontal cortex (PFC) function, yet the molecular underpinnings of this process remain poorly understood. We previously showed that SD disrupts sensorimotor gating by elevating prefrontal levels of the neurosteroid allopregnanolone (AP), a positive allosteric modulator of GABA-A receptors. Here we identify a complementary, mechanistically independent process whereby SD alters GABA-A currents in the PFC of mice and rats. SD reduced membrane expression of the chloride exporter KCC2, leading to intracellular chloride accumulation and a depolarizing shift in GABA-A receptor reversal potential that weakened GABAergic inhibition. Pharmacological normalization of chloride homeostasis with bumetanide fully rescued SD-induced deficits in sensorimotor gating and information encoding. SD also upregulated BDNF, and intra-PFC antagonism of its receptor TrkB restored KCC2 expression and normalized information processing, identifying BDNF–TrkB signaling as an upstream driver of chloride dysregulation. Notably, blocking AP synthesis rescued behavioral deficits without correcting chloride imbalance, confirming mechanistic independence. Finally, combined administration of AP and a KCC2 blocker produced information-processing deficits akin to those induced by SD. These findings identify TrkB-dependent disruption of prefrontal chloride homeostasis as a druggable mechanism underlying sleep loss–induced cognitive dysfunction.

## INTRODUCTION

Sleep is indispensable for cognitive performance, emotional regulation, and behavioral stability. Disruptions of sleep continuity and architecture are consistently associated with symptom severity across a broad spectrum of neuropsychiatric disorders (Benca et al., 1992; Frau et al., 2020). Although this clinical association is well established, the neurobiological mechanisms by which sleep loss destabilizes cortical information processing remain insufficiently characterized.

Sleep deprivation (SD) provides a robust experimental model to interrogate the behavioral and cognitive consequences of sleep loss. SD reliably impairs perception, sustained attention, and executive control (Killgore, 2010). In rodents, SD disrupts sensorimotor gating as measured by prepulse inhibition (PPI) (Cadeddu et al., 2022; Frau et al., 2008), a translational index of preattentive information filtering conserved across species (Braff et al., 2007). Consistent with these findings, PPI deficits have been reported in sleep-deprived humans (Petrovsky et al., 2014); but see (Vizeli et al., 2023) for conflicting results), underscoring the need for mechanistic clarification. Of note, the duration of SD required to produce reliable PPI impairment differs across species and strains: 24 hours is sufficient to disrupt gating in C57BL/6 mice (Cadeddu et al., 2022), whereas Sprague-Dawley rats require 72 hours of SD to produce comparable deficits (Frau et al., 2008).

Convergent evidence implicates the prefrontal cortex (PFC) as the key locus of SD-induced cognitive dysfunction (Krause et al., 2017). The PFC orchestrates executive control and top-down modulation of perception and behavior and critically contributes to sensorimotor gating (Swerdlow et al., 2001). Neuroimaging studies consistently demonstrate reduced PFC activity and disrupted functional connectivity with subcortical regions following SD (Verweij et al., 2014; Yang et al., 2018), paralleling impairments in attentional filtering, cognitive flexibility, and inhibitory control.

Our previous work has established neurosteroid signaling as a key modulator of SD-induced sensorimotor gating deficits. SD produces a marked elevation of allopregnanolone (AP) in the PFC, and this increase enables PPI impairments (Cadeddu et al., 2022; Frau et al., 2017). AP is a potent positive allosteric modulator of GABA-A receptors that enhances chloride conductance (Belelli & Lambert, 2005). Although AP can directly disrupt sensorimotor gating in C57BL/6 mice (Cadeddu et al., 2022), its effect on PPI in Sprague-Dawley rats depends on permissive conditions such as those produced by SD (Frau et al., 2017). This conditional relationship suggests that the behavioral impact of neurosteroid elevation is gated by additional factors and raises the question of whether upstream alterations in chloride homeostasis render GABAergic transmission vulnerable to neurosteroid potentiation. In the mature brain, the inhibitory efficacy of GABA-A receptors depends on low intracellular chloride concentrations, maintained by the coordinated activity of the cation–chloride cotransporters NKCC1, which imports chloride, and KCC2, which extrudes it (Blaesse et al., 2009). Perturbations of this balance, as observed during early development or after neuronal injury, result in intracellular chloride accumulation and depolarizing shifts in the GABA-A receptor reversal potential (E_GABA), thereby weakening or even reversing inhibitory transmission (Ben-Ari, 2002; Nabekura et al., 2002).

Recent studies have demonstrated that intracellular chloride levels in the mouse cortex exhibit diurnal fluctuations, with lower concentrations during the rest phase and higher concentrations during the active phase (Alfonsa et al., 2023; Pracucci et al., 2023). This rhythm appears to be partly governed by circadian clock mechanisms (Stangherlin et al., 2021). While maintaining wakefulness beyond the normal dark period does not abolish the physiological daytime decline in intracellular chloride (Pracucci et al., 2023), more robust SD protocols induce chloride accumulation in pyramidal neurons of the sensorimotor cortex (Alfonsa et al., 2023). These observations raise the possibility that chloride dysregulation represents a cellular mechanism linking SD to cortical dysfunction. However, it remains unresolved whether analogous processes occur in the PFC, a region critically implicated in the cognitive sequelae of sleep loss, and how they interact with neurosteroid elevations.

On this basis, we integrated behavioral, molecular, electrophysiological, two-photon chloride-imaging, and pharmacological approaches to test whether SD-induced information-processing deficits arise from disrupted chloride homeostasis in the PFC and whether elevated neurosteroid tone functions as a permissive factor when chloride gradients are compromised. To assess generalizability across species, we conducted these experiments in both mice and rats, which differ in their sensitivity to AP-induced gating deficits and in the duration of SD required to impair PPI. We examined the upstream mechanisms governing chloride transporter alterations and dissected the functional relationship between altered chloride gradients and neurosteroid signaling through targeted manipulation of AP synthesis and GABA-A receptor function.

## RESULTS

The following descriptions provide a summary of the results. Detailed statistical analyses for all experiments are provided in Supplementary Results.

### SD reduces KCC2 membrane expression in the prefrontal cortex

To assess whether SD alters chloride homeostasis in the PFC, we examined expression and phosphorylation of NKCC1 and KCC2 (**Figs. 1** and **S1-2**). SD was induced using the small-platform-over-water method (72 h in Sprague-Dawley rats; 24 h in C57BL/6 mice; the same durations required to produce PPI deficits). In both male mice and rats, SD significantly reduced membrane-associated KCC2 without affecting total protein levels (**Figs. 1B,E** and **S1B**), indicating impaired membrane trafficking rather than decreased synthesis. This reduction was accompanied by increased phosphorylation at threonine 1007 (**Figs. 1C,F**), a modification that promotes KCC2 internalization and functional inhibition (Friedel et al., 2015; Rinehart et al., 2009), resulting in an elevated pThr1007/total KCC2 ratio (**Fig. 1G**). In male rats, membrane and total NKCC1 expression were unchanged (**Figs. 1D** and **S1A**); however, phosphorylation at threonine 203/207/212 was increased (**Fig. S1C**), consistent with enhanced chloride import (Flemmer et al., 2002). These effects were independent of circadian phase (**Fig. S2**).

**Figure 1:**
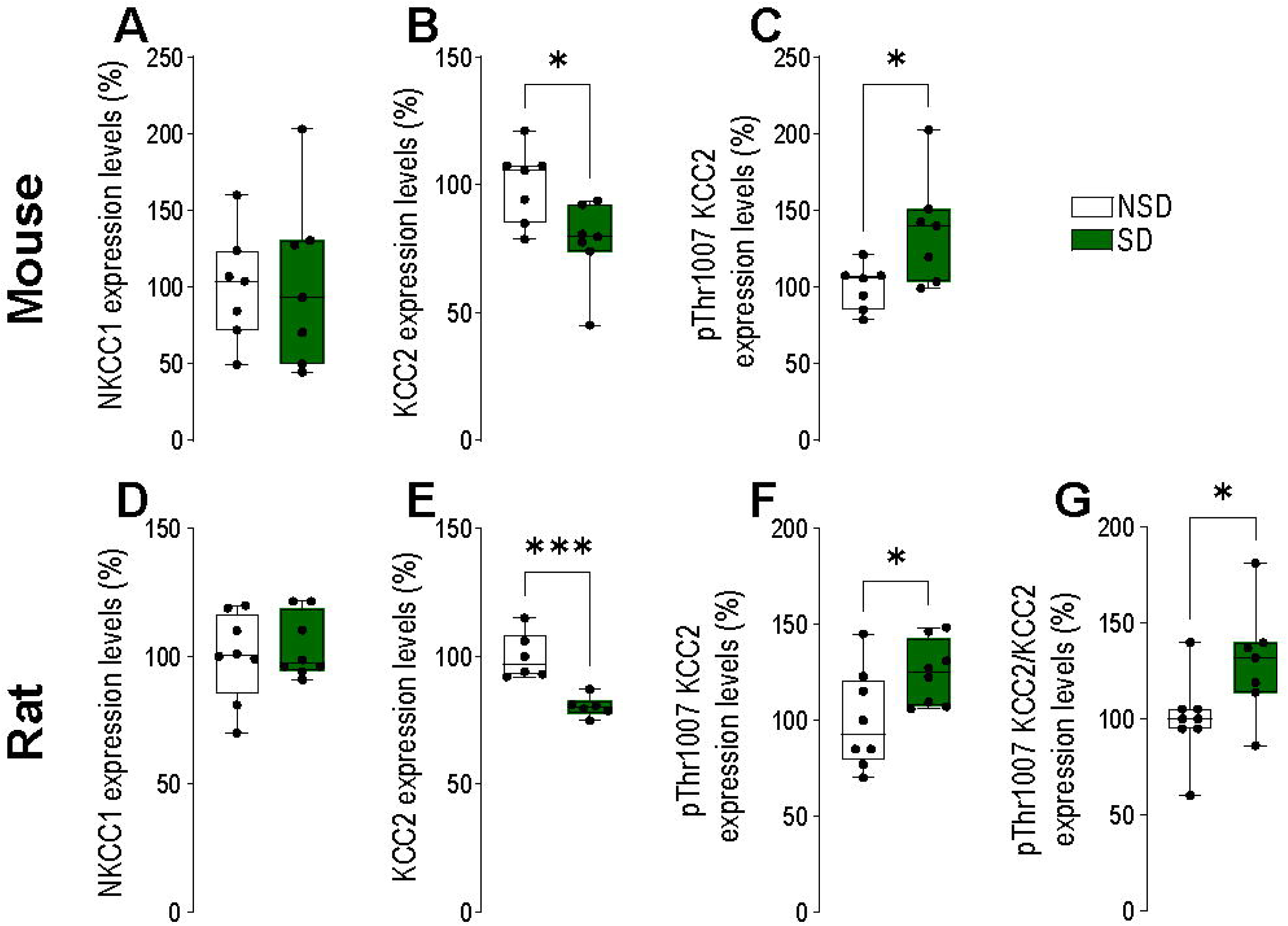
Sleep deprivation alters chloride transporter phosphorylation and membrane expression in the medial prefrontal cortex. Immediately following sleep deprivation (SD), male mice (A–C) and male rats (D–G) were sacrificed, and brain tissue was collected and flash-frozen. Western blot analyses of membrane extracts showed that in male mice, SD did not alter membrane expression of NKCC1 (A), but significantly reduced KCC2 levels (B) and increased its phosphorylation (C). Similar effects were observed in male rats exposed to SD (D–F) and were further confirmed by immunoprecipitation using an anti-pThr1007 KCC2 antibody (G). Data were analyzed using unpaired t tests and are presented as mean ± SEM, expressed as a percentage of non–sleep-deprived (NSD) controls. Each dot represents an individual animal. *p < 0.05, ***p < 0.001 for comparisons indicated by brackets.

### SD elevates intracellular chloride and depolarizes GABA-A signaling

The combined reduction in membrane KCC2 and increase in NKCC1 phosphorylation predicted intracellular chloride accumulation. *In vivo* two-photon imaging with LSSmClopHensor confirmed a marked elevation of intracellular chloride concentration ([Cl⁻]□) in medial PFC pyramidal neurons of male and female mice following SD. Mean [Cl⁻]□ increased from 13.6 mM in non–sleep-deprived controls, consistent with prior measurements at ZT5 (Pracucci et al., 2023), to 35.9 mM after SD (**Figs. 2A-C**), without detectable changes in intracellular pH. To assess the electrophysiological consequences of this shift in chloride homeostasis, gramicidin-perforated patch-clamp recordings were performed in medial PFC pyramidal neurons from male rats. These experiments revealed a depolarizing shift in E_GABA (**Figs. 2D–E**), indicating a reduced inhibitory driving force. SD also increased intrinsic excitability of pyramidal neurons, as evidenced by elevated firing frequency, reduced rheobase, and a hyperpolarized action potential threshold (**Figs. 2F-I**). In addition, SD reduced the paired-pulse ratio of evoked inhibitory postsynaptic currents (**Fig. 2J**). Together, these findings indicate that SD disrupts chloride homeostasis in the PFC, leading to depolarized GABA-A signaling and increased pyramidal neuron excitability.

**Figure 2.**
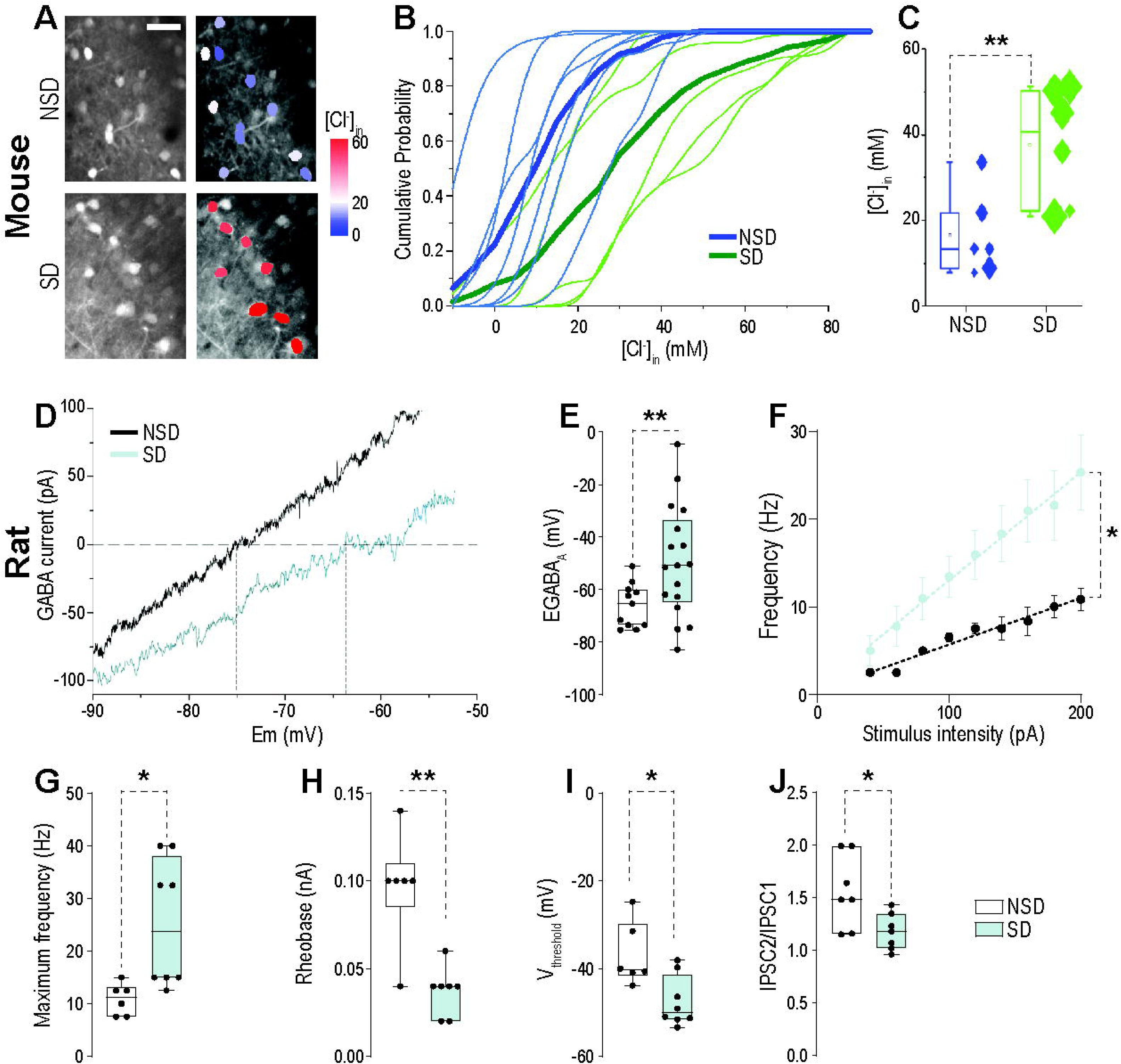
Sleep deprivation disrupts intracellular chloride homeostasis and increases excitability of prelimbic mPFC pyramidal neurons. Representative two-photon images (A) and quantification (B-C) of intracellular chloride concentration ([Cl⁻]□) measured using LSSmClopHensor in layer 2/3 pyramidal neurons of non-sleep-deprived (NSD, n = 6 mice, 560 cells) and sleep-deprived (SD, n = 6 mice, 308 cells) animals. In B, cumulative distributions of [Cl⁻]□ are shown for NSD (thin green lines) and SD mice (thin blue lines), with thick lines representing group averages. In C, each symbol represents the median [Cl⁻]□ for an individual mouse, and symbol size corresponds to twice the standard deviation of [Cl⁻]□ within that mouse. (D–J) Electrophysiological recordings from rats. (D) Current-voltage (I-V) curves of GABA-A receptor-mediated currents recorded from layer II-III prelimbic mPFC pyramidal neurons of NSD and SD rats using gramicidin-perforated patch-clamp technique. The reversal potentials (E_rev) correspond to the equilibrium potential of GABA-A currents (E_GABA, NSD, n = 9 neurons from 4 animals; SD, n = 7 neurons from 3 animals). (E) E_GABA measured for individual neurons in NSD (n = 9 neurons, 4 animals) and SD groups (n = 7 neurons, 3 animals), illustrating a depolarizing shift following SD. Each dot represents a single neuron. (F) SD pyramidal cells (n = 8 neurons, 4 animals) exhibit increased firing frequency in response to somatic current injections compared with NSD neurons (n = 6 neurons, 3 animals). (G) Quantification of maximum firing frequency in response to the largest current injection (200 pA). (H) Rheobase (minimal current intensity to evoke action potential) is lower in SD rats. (I) Voltage threshold for action potential initiation is reduced in SD rats. (J) Paired-pulse ratio (IPSC₂/IPSC₁) of GABA-A receptor-mediated IPSCs is reduced in SD rats. *p < 0.05, **p < 0.01, ***p < 0.001 for comparisons indicated by brackets. Unless otherwise indicated, each dot represents an individual animal.

### Restoration of chloride homeostasis rescues SD-induced deficits in information processing

We therefore assessed whether chloride dysregulation is causally required for the cognitive impairments elicited by SD. In male and female mice, systemic administration of the NKCC1 inhibitor bumetanide dose-dependently reversed SD-induced deficits in prepulse inhibition (PPI), without affecting baseline startle reactivity (**Figs. 3A–D**). Similar results were obtained in male rats, in which bumetanide fully restored PPI following SD (**Figs. 3E-F**). Because PPI primarily indexes preattentional filtering, we next examined whether chloride modulation could also mitigate SD-induced impairments in information encoding. To this end, we tested the impact of bumetanide during the encoding phase of a novel object recognition (NOR) task in SD-exposed rats (**Fig. 3G**). Immediately after SD, animals were exposed to two identical objects and subsequently returned to their home cages for 24 h to allow sleep recovery. During the retrieval phase, rats encountered one familiar and one novel object. SD markedly reduced the novel exploration index (NEI), consistent with impaired encoding, and bumetanide significantly rescued NEI in SD rats.

**Figure 3.**
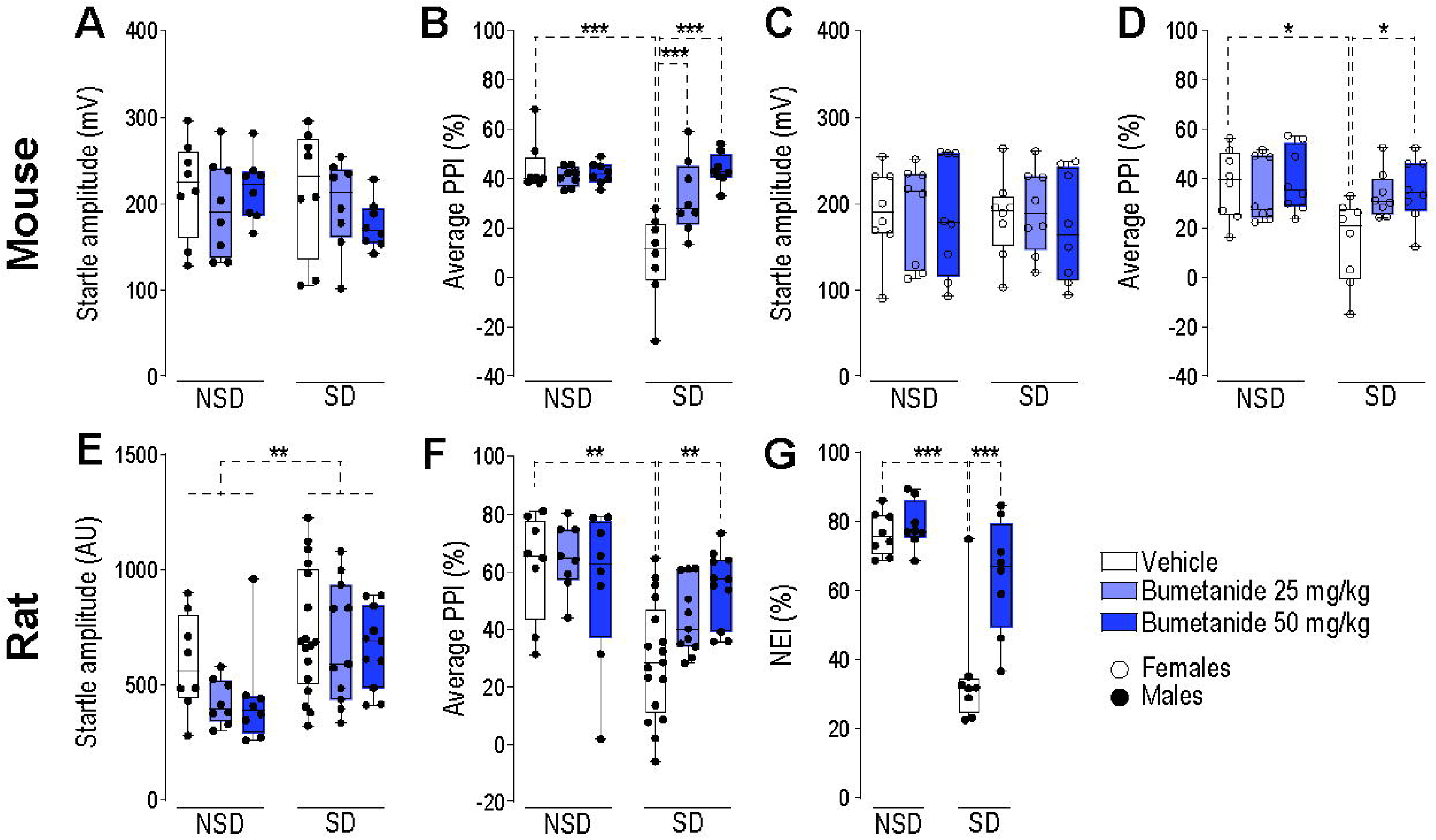
The NKCC1 inhibitor bumetanide rescues sleep deprivation-induced deficits in prepulse inhibition and information encoding. Effects of bumetanide on startle reflex amplitude (A, C, E), prepulse inhibition (PPI; B, D, F), and novel object recognition (novelty exploration index, %NEI; G) following systemic administration of bumetanide (25–50 mg/kg, IP). (A–B) Male mice: bumetanide had no effect on startle amplitude (A) but dose-dependently rescued the SD-induced PPI deficit (B). (C–D) Female mice: bumetanide had no effect on startle amplitude (C) but dose-dependently rescued the SD-induced PPI deficit (D). (E–G) Male rats: SD increased startle amplitude, which was not affected by bumetanide administration (E); bumetanide dose-dependently reversed the SD-induced PPI deficit (F) and normalized SD-induced deficits in memory encoding as assessed by the novelty exploration index (G). Data are presented as mean ± SEM and were analyzed using two-way ANOVA. *p < 0.05, **p < 0.01, ***p < 0.001 for comparisons indicated by brackets. Each dot represents an individual animal.

### SD upregulates BDNF and GABA-A receptor α4 subunits in the PFC

We next investigated whether SD alters GABA-A receptor expression and subunit composition. In male rats, SD selectively increased expression of the GABA-A α4 subunit without altering α1 or δ levels (**Figs. 4A-C**). Because α4-containing receptors exhibit heightened neurosteroid sensitivity (Smith et al., 2009; Stell et al., 2003), this finding suggests a mechanism by which SD amplifies the impact of elevated AP.

**Figure 4.**
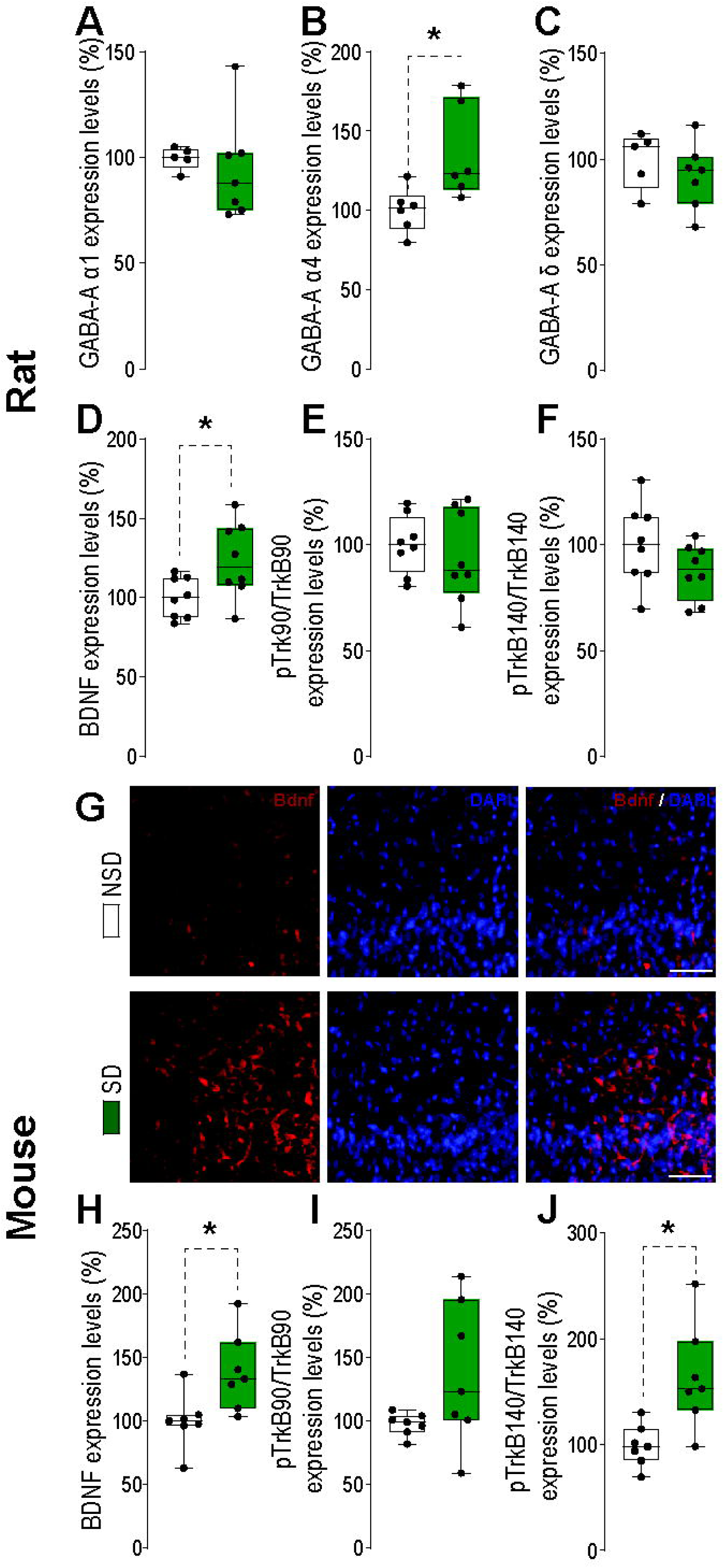
Sleep deprivation selectively alters GABA-A receptor subunit expression and BDNF signaling in the medial prefrontal cortex. (A–F) Male rats. Immediately following sleep deprivation (SD), rats were sacrificed and brain tissue was collected and frozen. Western blot analyses were performed to assess GABA-A receptor subunits (A–C) and components of the BDNF signaling pathway (D–F). SD did not affect expression of the α1 (A) or δ (C) subunits but significantly increased the expression of the α4 subunit (B). SD increased BDNF protein levels (D) without altering phosphorylation of the TrkB receptor, either in its truncated (TrkB90; E) or full-length (TrkB140; F) forms. (G–J) Male mice. Analyses revealed increased BDNF protein levels (G–H), with panel G presenting representative immunofluorescence images and panel H showing the quantification of Western blot analyses. This increase was accompanied by enhanced phosphorylation of full-length TrkB140 (J), whereas no changes were detected in the truncated TrkB90 form (I). Data are presented as mean ± SEM, expressed as a percentage of non–sleep-deprived (NSD) controls, and were analyzed using unpaired t-tests. *p < 0.05 for comparisons indicated by brackets. Each dot represents an individual animal.

We next sought to identify the upstream mechanism responsible for both α4 upregulation and membrane KCC2 downregulation. BDNF emerged as a compelling candidate, given its established role in regulating GABA-A receptor subunit expression (Roberts et al., 2006) and KCC2 membrane trafficking (Rivera et al., 2002), as well as prior evidence that SD elevates BDNF levels (Giacobbo et al., 2016; Schmitt et al., 2016). Consistent with this hypothesis, SD produced a significant increase in BDNF protein expression in the mPFC of both male rats (**Fig. 4D**) and male mice (**Figs. 4G-H**), without altering proBDNF levels (**Figs. S3A-D** and **S4A**). Similar increases in BDNF levels were also observed in the nucleus accumbens and amygdala of male rats (**Figs. S3E-G**). In addition, SD modulated TrkB receptor signaling with species-specific differences in the activation of downstream pathways. Total TrkB expression remained unchanged in both mice and rats (**Figs. S3H-I** and **S4B-C**). In male rats, TrkB phosphorylation was not altered (**Figs. 4E–F**). In contrast, male mice showed increased phosphorylation of the full-length TrkB140 isoform (**Figs. 4I–J**), consistent with enhanced BDNF signaling activity.

### TrkB blockade prevents SD-induced molecular and behavioral deficits

To test whether TrkB signaling plays a causal role in the information-processing deficits caused by SD, we administered the TrkB antagonist ANA-12 to male rats either systemically (**Figs. 5A-B**) or locally into the mPFC (**Figs. 5C-I**). Systemic ANA-12 partially rescued PPI deficits (**Fig. 5B**). Furthermore, intra-PFC ANA-12 fully rescued SD-induced deficits in PPI and information encoding in NOR (**Fig.5D-E**). ANA-12 normalized SD-induced alterations in inhibitory signaling, restoring GABA-A receptor subunit composition (**Figs. 5F-G**) and chloride transporter regulation (**Figs. 5H-I** and **S5**). Taken together, these data indicate that TrkB signaling is necessary for the coordinated changes in receptor composition and chloride homeostasis underlying SD-induced information-processing impairments.

**Figure 5.**
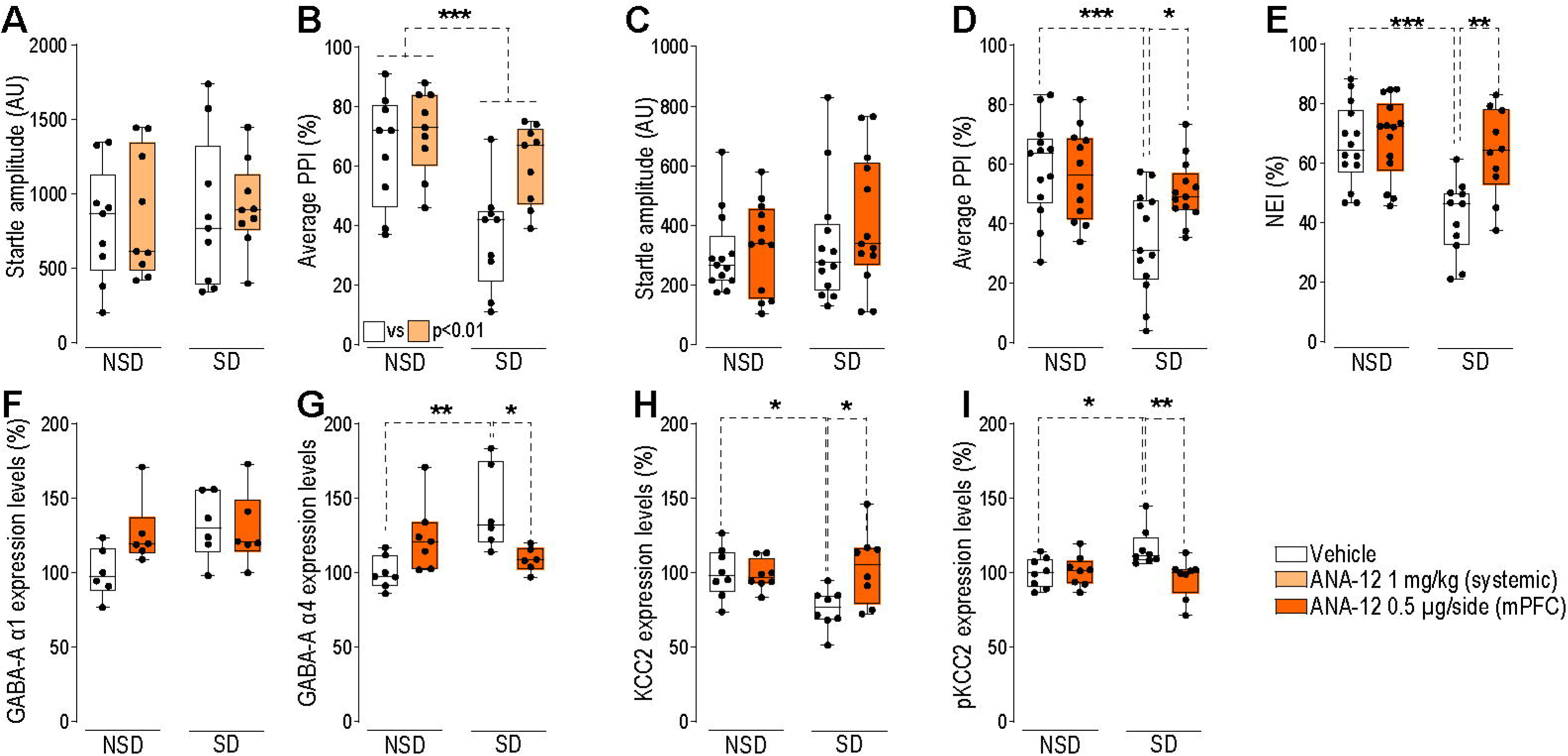
The TrkB antagonist ANA-12 rescues sleep deprivation-induced deficits in prepulse inhibition and information encoding. Effects of ANA-12 on startle reflex amplitude (A, C), prepulse inhibition (PPI; B, D), and novel object recognition (E) in sleep-deprived (SD) and non–sleep-deprived (NSD) male rats following systemic (1 mg/kg, IP; A,B) or intra-prefrontal cortex (PFC; 0.5 µg/side; C–J) administration of ANA-12. Systemic ANA-12 had no effect on startle amplitude but partially rescued the SD-induced PPI deficit (B). Intra-PFC administration of ANA-12 did not alter startle amplitude (C) and fully reversed the SD-induced PPI deficit (D). ANA-12 also normalized SD-induced deficits in memory encoding (E), as assessed by the novelty exploration index (NEI). In addition, ANA-12 did not affect GABA-A α1 subunit expression (F) but restored SD-related alterations in GABA-A α4 subunit (G), membrane KCC2 levels (H), and KCC2 phosphorylation (I) in SD rats. Data are presented as mean ± SEM and were analyzed using two-way ANOVA. *p < 0.05, **p < 0.01, ***p < 0.001 for comparisons indicated by brackets. Each dot represents an individual animal.

### Neurosteroid and chloride pathways operate independently

We next examined whether neurosteroid elevation and chloride dysregulation represent independent pathogenic mechanisms. In line with previous results (Frau et al., 2017), the 5α-reductase inhibitor finasteride, which blocks AP biosynthesis, fully rescued PPI deficits induced by SD (**Fig. 6**). In male and female mice, systemic finasteride (12.5–25 mg/kg) rescued SD-induced PPI deficits without affecting startle reactivity (**Figs. 6A-D**). In male rats, intra-PFC finasteride (0.5 µg/side) similarly rescued PPI deficits (**Figs. 6F-G**) and normalized NEI (**Fig. 6H**). Strikingly, despite complete behavioral rescue, finasteride failed to correct the underlying molecular disruptions. In SD male and female mice, intracellular chloride remained elevated after finasteride treatment (**Fig. 6E**). In male rats, membrane KCC2 expression was not restored (**Fig. 6J**), GABA-A α4 upregulation persisted (**Figs. S6A–C**), and BDNF levels were unchanged (**Figs. S6D,E**). These dissociations establish that AP elevation and chloride dysregulation are mechanistically independent pathways that each contribute to SD-induced deficits.

**Figure 6.**
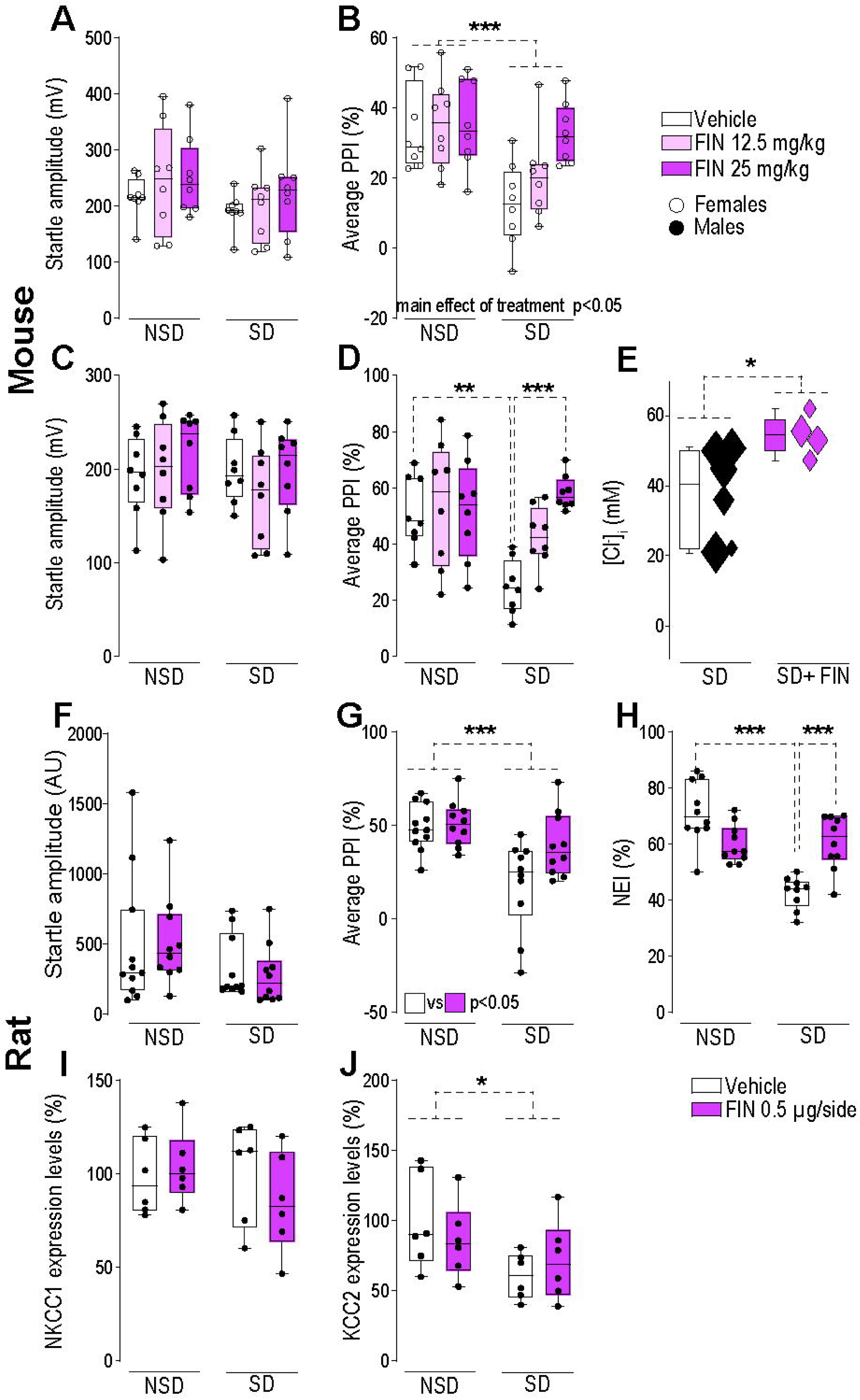
The 5α-reductase inhibitor finasteride rescues sleep deprivation-induced deficits in prepulse inhibition and information encoding without restoring chloride homeostasis. (A–E) Male and female mice. Effects of finasteride (FIN; 12.5–25 mg/kg, IP) on startle reflex amplitude (A, C), prepulse inhibition (PPI; B, D), and intracellular chloride concentration (E) in sleep-deprived (SD) and non–sleep-deprived (NSD) mice. FIN had no effect on startle amplitude in either female (A) or male (C) mice. FIN produced a main effect on PPI in females (B) and dose-dependently rescued the SD-induced PPI deficit in males (D). The control level of intracellular chloride concentration [Cl⁻] in SD mice was not rescued by FIN injection (before FIN: n = 6 mice, 308 cells; after FIN: n = 4 mice, 119 cells); E). (F–J) Male rats. Effects of intra-PFC FIN (0.5 µg/side) on startle amplitude (F), PPI (G), novel object recognition (H), and chloride transporter expression (I,J). FIN did not affect startle amplitude (F) but reversed the SD-induced PPI deficit (G) and normalized SD-induced deficits in memory encoding as assessed by the novelty exploration index (H). FIN produced no changes in NKCC1 (I) or KCC2 (J) expression levels. Data are presented as mean ± SEM and were analyzed using two-way ANOVA. *p < 0.05, **p < 0.01, ***p < 0.001 for comparisons indicated by brackets. Each dot represents an individual animal.

### Intracellular chloride accumulation synergizes with AP to produce gating deficits

Finally, we tested whether elevated AP and disrupted chloride homeostasis are sufficient in combination to impair sensorimotor gating. In male rats, neither AP nor the selective KCC2 inhibitor VU0463271 (Delpire et al., 2009) alone impaired PPI (**Fig. 7**). However, their combined administration produced robust PPI deficits comparable to those observed in SD animals (**Figs. 7B,D**). These findings indicate that the combined elevation of AP and disruption of chloride homeostasis is sufficient to recapitulate the SD-induced gating deficit.

**Figure 7.**
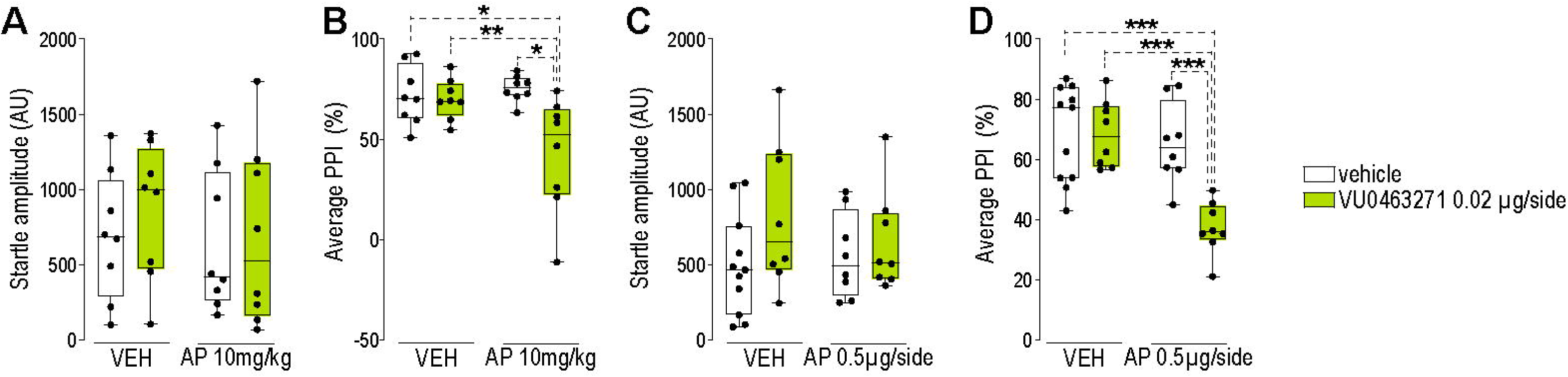
Combined administration of allopregnanolone and the KCC2 inhibitor VU0463271 reproduces sleep deprivation-like deficits in prepulse inhibition. Effects of allopregnanolone (AP) and VU0463271 on startle reflex amplitude (A, C) and prepulse inhibition (PPI; B, D) in non–sleep-deprived male rats. AP was administered either systemically (10 mg/kg, IP; A,B) or intra-prefrontal cortex (PFC; 0.5 µg/side; C,D), whereas VU0463271 was administered intra-PFC (0.02 µg/side) in all conditions. Startle amplitude was not affected by AP, VU0463271, or their combination under either administration paradigm (A, C). In contrast, combined administration of AP and VU0463271 significantly reduced PPI, whereas neither compound alone altered PPI (B, D). Data are presented as mean ± SEM and were analyzed using two-way ANOVA. *p < 0.05, **p < 0.01, ***p < 0.001 for comparisons indicated by brackets. Each dot represents an individual animal.

## DISCUSSION

The findings of this study demonstrate that SD impairs sensorimotor gating and information encoding in rodents through disruption of chloride homeostasis in the prefrontal cortex. SD reduced membrane expression and functional activity of the chloride extruder KCC2 via increased phosphorylation at Thr1007, elevated intracellular chloride concentrations, and induced a depolarizing shift in E_GABA. These ionic disturbances were accompanied by enhanced intrinsic neuronal excitability and selective upregulation of the neurosteroid-sensitive GABA-A α4 subunit. Bumetanide, which restores chloride gradients through inhibition of NKCC1, fully reversed the SD-induced deficits in sensorimotor gating and information encoding. Local blockade of TrkB in the medial prefrontal cortex normalized both the molecular alterations and the associated behavioral impairments, indicating that TrkB signaling is required for KCC2 downregulation following SD. Notably, finasteride restored behavioral performance without correcting chloride dysregulation, whereas combined administration of AP and a KCC2 inhibitor reproduced the information-processing deficits of SD in non–sleep-deprived animals. Collectively, these findings establish chloride dysregulation as a central mechanism that converges with neurosteroid elevation to produce SD-induced information-encoding deficits.

Recent studies have shown that intracellular chloride concentrations exhibit diurnal fluctuations in the mouse cortex, with lower levels during the rest phase and higher levels during the active phase (Alberio et al., 2025; Alfonsa et al., 2023; Pracucci et al., 2023). This rhythm appears to be circadian in origin and remains stable under conditions of mild sleep perturbation (Pracucci et al., 2023; Stangherlin et al., 2021). In contrast, more prolonged or severe SD protocols increase intracellular chloride in cortical pyramidal neurons (Alfonsa et al., 2023). The present study extends these observations to the prefrontal cortex and delineates distinct mechanistic substrates. Whereas brief extensions of wakefulness primarily influenced NKCC1 function, prolonged SD lasting 24 to 72 hours reduced membrane-associated KCC2 without altering total protein abundance, indicating a trafficking deficit rather than transcriptional downregulation. Although NKCC1 membrane levels were unchanged, SD increased phosphorylation at Thr203/207/212, consistent with enhanced chloride influx. These findings indicate that KCC2 downregulation constitutes the principal driver of chloride accumulation in the prefrontal cortex following prolonged SD, with NKCC1 hyperphosphorylation contributing in a modulatory capacity. Minor discrepancies with earlier reports (Alfonsa et al., 2023) likely reflect regional heterogeneity, differences in the duration of SD, or both. Systematic analysis of the temporal dynamics of chloride regulation during SD will therefore be essential.

A major mechanistic contribution of this study is that BDNF–TrkB signaling is required for SD-induced downregulation of KCC2. SD is a well-established stimulus for BDNF upregulation in cortical neurons (Cirelli et al., 2004; Cirelli & Tononi, 2000; Faraguna et al., 2008; Huber et al., 2007), and consistent with this, we observed significant increases in mature BDNF protein in the PFC of both rats and mice following SD. However, TrkB receptor phosphorylation exhibited species-specific patterns: in male mice, SD increased phosphorylation of full-length TrkB140, whereas in male rats, TrkB phosphorylation remained unchanged despite elevated BDNF levels. This discrepancy may reflect differences in the temporal dynamics of TrkB activation, such that peak phosphorylation occurred outside the sampling window in rats, or the differing SD durations (24 hours in mice vs. 72 hours in rats), which may engage distinct phases of the BDNF–TrkB signaling cascade. Critically, despite these differences, intra-mPFC administration of the TrkB antagonist ANA-12 in rats fully reversed both the behavioral and molecular consequences of SD, restoring KCC2 expression, normalizing its phosphorylation state, and correcting GABA-A α4 subunit upregulation. This pharmacological rescue demonstrates that ongoing TrkB signaling is required for SD-induced chloride dysregulation in rats and suggests that a similar mechanism operates in mice, given the conserved BDNF upregulation and KCC2 downregulation observed across both species. Taken together, these findings are consistent with an extensive literature linking BDNF–TrkB signaling to KCC2 regulation (Puskarjov et al., 2015; Rivera et al., 2002; Wardle & Poo, 2003) and position this pathway as a conserved mechanistic driver of GABAergic dysfunction following sleep loss.

Although the present experiments did not directly dissect the mechanisms underlying BDNF upregulation during SD, several candidate processes merit consideration. Prolonged wakefulness increases extracellular glutamate concentrations and enhances NMDA and AMPA receptor activation, thereby promoting BDNF transcription from activity-dependent promoters (Bessho et al., 1993; Tao et al., 1998; Zheng et al., 2012). Concurrently, SD induces neuroinflammatory responses characterized by elevated proinflammatory cytokines (Zielinski et al., 2016), which may stimulate BDNF release from astrocytes (Saha et al., 2006) and microglia (Trang et al., 2009). Robust microglial activation has been documented in the prefrontal cortex during SD (Bellesi et al., 2017; Wadhwa et al., 2017), raising the possibility that microglia-to-neuron BDNF signaling contributes to chloride dysregulation under conditions of sleep loss. This mechanism was originally characterized in neuropathic pain models (Coull et al., 2005; Ferrini & De Koninck, 2013), but the shared involvement of BDNF–TrkB–KCC2 signaling suggests it may generalize to other contexts of sustained neuronal challenge. This hypothesis requires direct experimental evaluation.

Another central finding of this work is that chloride dysregulation, although necessary, is not by itself sufficient to produce the information-processing deficits associated with SD; behavioral impairment arises from convergence with elevated neurosteroid signaling. Finasteride completely reversed SD-induced deficits in prepulse inhibition and novel object recognition, yet did not normalize intracellular chloride levels, membrane KCC2 expression, or GABA-A α4 subunit upregulation. These dissociations demonstrate that chloride dysregulation and AP elevation represent mechanistically distinct pathways whose interaction is required for behavioral impairment. This interpretation is strengthened by our finding that neither AP administration nor KCC2 inhibition alone disrupted sensorimotor gating in non–sleep-deprived animals, whereas their combination fully reproduced the deficits observed after SD.

This dual-pathway model provides a coherent explanation for the state-dependent behavioral effects of AP. GABA-A receptor–mediated currents are determined by the interaction between channel conductance and electrochemical driving force. Under physiological chloride gradients, increased conductance induced by AP enhances inhibition without destabilizing network dynamics. When chloride gradients are depolarized but AP levels remain within normal range, GABA conductance is insufficient to generate pathological depolarizing currents. In contrast, when elevated AP increases GABA-A conductance in the presence of a depolarized E_GABA, large depolarizing GABA currents can emerge, compromising inhibitory precision and network timing. Upregulation of α4-containing, extrasynaptic GABA-A receptors, which exhibit high neurosteroid sensitivity, likely amplifies this interaction. This conductance-by-driving-force framework also accounts for the ability of finasteride to rescue behavior despite persistent chloride dysregulation: by reducing conductance, it limits depolarizing GABA currents even when E_GABA remains shifted.

An important conceptual issue concerns whether AP elevation during SD is maladaptive or compensatory. Neurosteroid synthesis may confer neuroprotection by attenuating hyperexcitability, oxidative stress, and neuroinflammation associated with prolonged wakefulness. From this standpoint, cognitive impairment may represent a trade-off in which strengthened GABAergic tone stabilizes neuronal activity at the expense of the temporal precision required for information encoding. This framework generates a testable prediction: if AP elevation serves a protective function, then chronic blockade of neurosteroid synthesis during SD should unmask excitotoxic damage or exacerbate neuroinflammation, even as it preserves information-processing performance. This possibility warrants direct empirical evaluation.

The mechanisms delineated here may have relevance for neuropsychiatric disorders characterized by sleep disruption and deficits in information processing. Mania provides a particularly salient parallel: SD can precipitate manic episodes (Colombo et al., 1999), and mania is associated with marked impairments in sensorimotor gating (Perry et al., 2001). Chloride dysregulation, potentially interacting with altered neurosteroid signaling, may therefore contribute to affective switching in bipolar disorder. Similar reasoning may apply to schizophrenia, in which PPI deficits are a hallmark feature and sleep disturbances are pervasive yet mechanistically underexplored. More broadly, any condition in which chronic sleep disruption co-occurs with GABAergic dysfunction could engage the dual-pathway mechanism described here, suggesting that chloride homeostasis deserves closer scrutiny as a transdiagnostic substrate.

Several limitations should be considered. Although we employed the widely validated small-platform method to induce SD, this approach does not completely dissociate sleep loss from mild stress exposure. While prolonged wakefulness is intrinsically stressful, several observations argue against stress as the sole determinant of the observed phenotype. SD-induced alterations in KCC2 and NKCC1 phosphorylation occurred independently of circadian phase, indicating that the molecular changes cannot be attributed simply to time-of-day effects or acute stress responses. Furthermore, the specificity of TrkB blockade in rescuing both molecular and behavioral phenotypes supports involvement of a defined BDNF-dependent pathway rather than a nonspecific stress mechanism. Prior work from our group also demonstrated that placement on a larger platform, which permits sleep while maintaining comparable confinement conditions, did not induce PPI deficits (Frau et al., 2017), further dissociating the behavioral phenotype from generalized stress effects. Nevertheless, a contributory role of stress cannot be entirely excluded, and future studies employing alternative SD paradigms or direct stress manipulations will be required to clarify this issue.

Another limitation concerns sex as a biological variable. Both male and female mice were included in chloride imaging and behavioral pharmacology experiments and exhibited qualitatively similar responses, providing preliminary evidence that these mechanisms are not sex-specific. However, the study was not adequately powered to support formal sex-stratified analyses. Given well-documented sex differences in neurosteroid signaling, stress responsivity, and sleep architecture (Hodes et al., 2024; Raciti et al., 2023; Schwarz & Schiza, 2024), future studies specifically designed to interrogate sex as a biological variable will be necessary to determine whether the mechanisms described here operate equivalently across sexes or exhibit meaningful dimorphism.

In summary, this study identifies a previously unrecognized mechanism by which SD disrupts prefrontal cortical function and information encoding through TrkB-dependent downregulation of KCC2 and consequent intracellular chloride accumulation. Critically, chloride dysregulation alone is not sufficient to impair information processing; behavioral deficits emerge from its convergence with elevated neurosteroid signaling, establishing a dual-pathway model in which the conductance and driving-force determinants of GABA-A currents are independently compromised. These findings refine our understanding of the neurobiological consequences of SD and identify chloride homeostasis as a mechanistically defined, pharmacologically accessible target for cognitive impairment associated with sleep loss and neuropsychiatric conditions marked by disturbed sleep–wake regulation.

## METHODS

### Animals

Adult male and female C57BL/6J mice (8–12 weeks) and adult male Sprague Dawley rats (250–320 g) were housed under standard conditions (22°C, 12:12 light:dark cycle) with ad libitum access to food and water. All procedures were approved by the respective Institutional Animal Care and Use Committees at participating institutions and conducted in accordance with EU Directive 2010/63/EU and the NIH Guide for the Care and Use of Laboratory Animals.

### Sleep deprivation

SD was performed using the small-platform-over-water method (Cadeddu et al., 2022; Frau et al., 2008, 2017). Rats were sleep-deprived for 72 hours; mice for 24 hours. Control animals were housed individually under identical conditions. Full details are provided in Supplementary Methods.

### Pharmacological treatments

Drugs were administered systemically (IP) or by stereotaxic microinfusion into the medial PFC: allopregnanolone (10 mg/kg IP or 0.5 μg/0.5 μl/side intra-PFC), bumetanide (25-50 mg/kg IP), finasteride (12.5-25 mg/kg IP or 0.5 μg/0.5 μl/side intra-PFC), VU0463271 (0.02µg/0.5 μl/side intra-PFC), and ANA-12 (1 mg/kg IP or 0.5 μg/0.5 μl intra-PFC). Vehicle-treated animals served as controls. Drug sources and preparation details are provided in Supplementary Methods.

### Behavioral testing

Sensorimotor gating was evaluated using prepulse inhibition (PPI) of the acoustic startle reflex (a pre-attentive gating measure), as previously described (Frau et al., 2016). Startle stimuli (120 dB, 40 ms) were delivered either alone or preceded by prepulse stimuli (74, 78, or 82 dB). Percent PPI at each prepulse intensity was calculated as [(mean startle amplitude on pulse-alone trials − mean startle amplitude on prepulse+pulse trials) / mean startle amplitude on pulse-alone trials] × 100. Because no significant interactions between prepulse intensity and any experimental factors were detected, PPI values were averaged across the three prepulse levels for all analyses.

Novel object recognition (NOR) was conducted as described previously (Bortolato et al., 2010). Rats were first habituated to the testing arena. Following habituation, they underwent SD (72 h) and were then immediately subjected to the encoding phase, during which they were exposed to two identical objects. Rats were subsequently returned to their home cages and allowed ad libitum sleep for 24 h. Retrieval was then assessed by presenting one familiar and one novel object. The novelty exploration index (NEI) was calculated as [time exploring novel object / (time exploring novel + time exploring familiar)] × 100.

### In vivo chloride imaging

Intracellular chloride was measured using the genetically encoded ratiometric biosensor LSSmClopHensor, selectively expressed in cortical pyramidal neurons. Expression was achieved by viral transfection using a mixture of two viral vectors: AAV9-EF1a-DIO(LSSmClopHensor) and AAV9-CaMKII-Cre. Ratiometric imaging at five excitation wavelengths (800, 830, 860, 910, and 960 nm) provided pH-corrected chloride measurements as previously described (Pracucci et al., 2023). Absolute concentrations were determined by calibration with ionophores.

### Electrophysiology

Whole-cell patch-clamp recordings were performed in acute coronal slices (300 μm) containing the mPFC. Intrinsic excitability was assessed in current-clamp mode using depolarizing current steps. GABA-A receptor reversal potential (E_GABA) was measured using gramicidin perforated-patch recordings to preserve native intracellular chloride concentrations. IPSCs were recorded in voltage-clamp mode with ionotropic glutamate receptors blocked (AP5, CNQX). Full protocols are detailed in Supplementary Methods.

### Western blotting

PFC tissue was dissected and processed for total or membrane-enriched protein fractions. Proteins were separated by SDS-PAGE, transferred to nitrocellulose, and probed with antibodies against KCC2, NKCC1, phospho-KCC2 (Thr1007), phospho-NKCC1 (Thr203/207/212), GABA-A receptor subunits (α1, α4, δ), and BDNF. Band densities were normalized to β-actin or total protein. Antibody details and full protocols are provided in Supplementary Methods.

### Statistics

Data were analyzed using unpaired t-tests for two-group comparisons or multi-way ANOVA followed by Tukey’s post hoc tests for experiments with multiple factors. PPI data were collapsed across prepulse intensities as described above. Statistical significance was set at p < 0.05. Sample sizes were determined by power analysis based on prior studies (Frau et al., 2008, 2017). Full statistical details for each experiment are provided in Supplementary Methods and Results.

### Data availability

The data that support the findings of this study are available from the corresponding author upon reasonable request.

## ACKNOWLEDGEMENT

This study was partially supported by the NIH grant R56 MH130006, R01 MH104603, and R01 AA030256 (to MB) and by “Fondazione di Sardegna” (to RF, ID: F73C23001630007) and “Fondazione Guy Everett” (to RF). The authors wish to thank all the personnel of CESAST (Centro Servizi di Ateneo per gli Stabulari) for their skillful assistance in animal care. GN was supported by PRIN 2022°58M7F. GP was partially supported by Telethon grant GMR24T1119. NEXTGENERATIONEU (NGEU); the Ministry of University and Research; the National Recovery and Resilience Plan; project MNESYS (PE0000006)—A multiscale integrated approach to the study of the nervous system in health and disease (DN. 1553 11.10.2022) to GMR. SL was supported by PRIN 2022 2022YSTP5L_LS5 to SL.

## AUTHOR CONTRIBUTIONS

MB, GMR, and RF conceived the study. CB, GMR, GN, and MB supervised the analyses. CB, EC, FB, GB, GN, GP, LC, MM, MS, PF, SaS, SiS, SL, FT, and VS performed experiments and analyses. CB, EC, FB, GB, GN, GP, LC, MM, MS, PF, SaS, SiS, SL, FT, MP, and VS performed experiments and analyses. EC, FB, GB, GP, LC, SaS, SL, and VS acquired data. MB wrote the manuscript. CB, GMR, GN, GP, MB, MM, RF, SiS, and SL reviewed and edited the manuscript.

