## Supplementary information for "Sleep deprivation impairs information processing via dysregulation of chloride homeostasis in the prefrontal cortex"

### SUPPLEMENTARY FIGURES

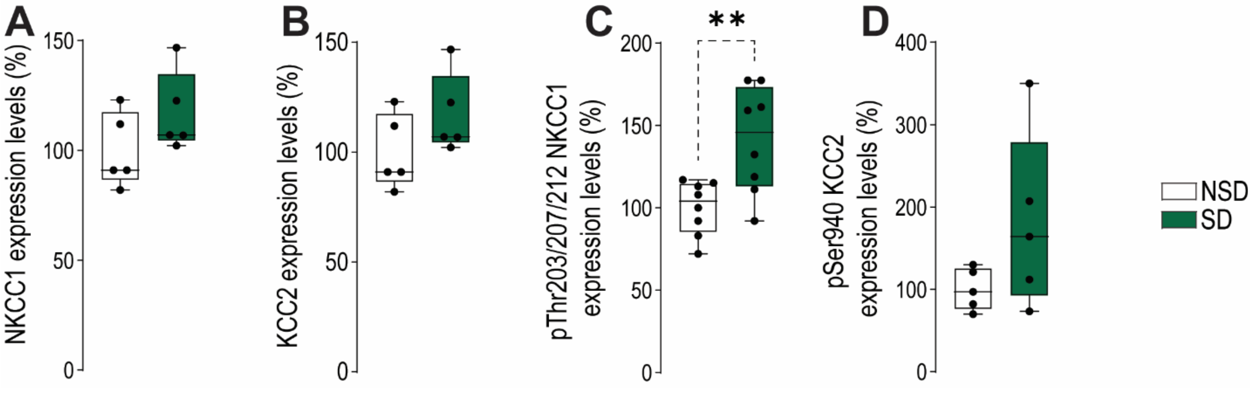


**Supplementary Figure 1 (Related to Figure 1).** Immediately after sleep deprivation (SD), rats were sacrificed and brain tissue was collected and flash-frozen. Western blot analyses of total lysates showed no changes in total NKCC1 (A) or KCC2 (B) expression. In contrast, analyses of membrane-enriched fractions revealed a significant increase in NKCC1 phosphorylation following SD compared to non–sleep-deprived (NSD) controls (C). Phosphorylation of KCC2 at Ser940 in membrane-enriched fractions was not altered (D). Data are presented as mean ± s.e.m. and were analyzed using unpaired two-tailed t-tests. ** $p < 0.01$ for the comparison indicated by brackets. Each dot represents one animal.

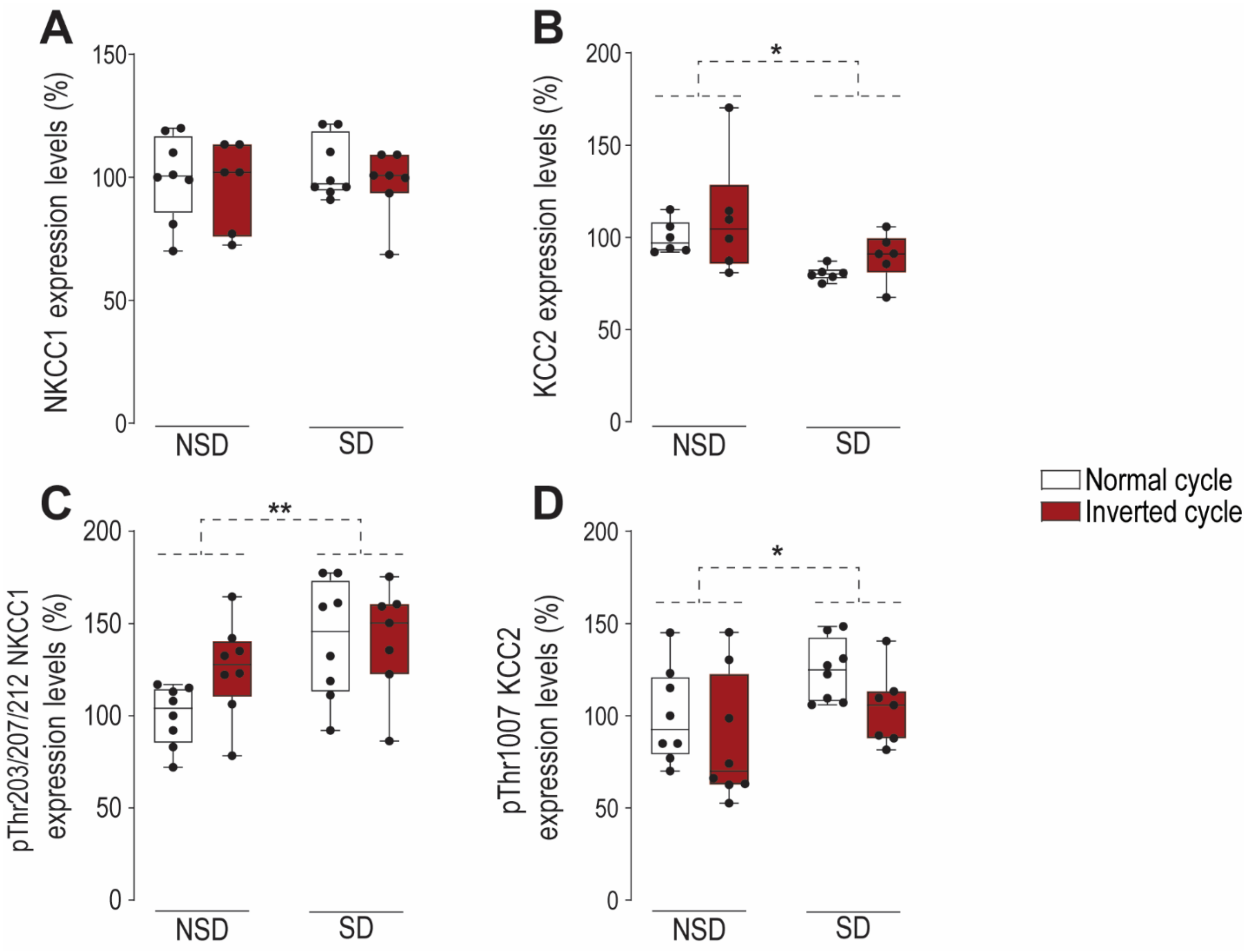


**Supplementary Figure 2 (Related to Figure 1). Sleep deprivation-induced alterations in chloride transporter expression and phosphorylation are independent of circadian phase.** Immediately following sleep deprivation during either the light or dark phase, rats were sacrificed and brain tissue was collected and flash-frozen. Western blot analyses were performed to assess membrane NKCC1 (A), membrane KCC2 (B), phosphorylated NKCC1 at Thr203/207/212 (C), and phosphorylated KCC2 at Thr1007 (D). Independent of circadian phase, SD significantly altered membrane KCC2 expression (B) and both NKCC1 (C) and KCC2 (D) phosphorylation in membrane-enriched fractions. Data are presented as mean ± SEM and were by two-way ANOVA.*, $p<0.05$, **, $p < 0.01$ for comparison indicated by brackets. Each dot represents an individual animal.

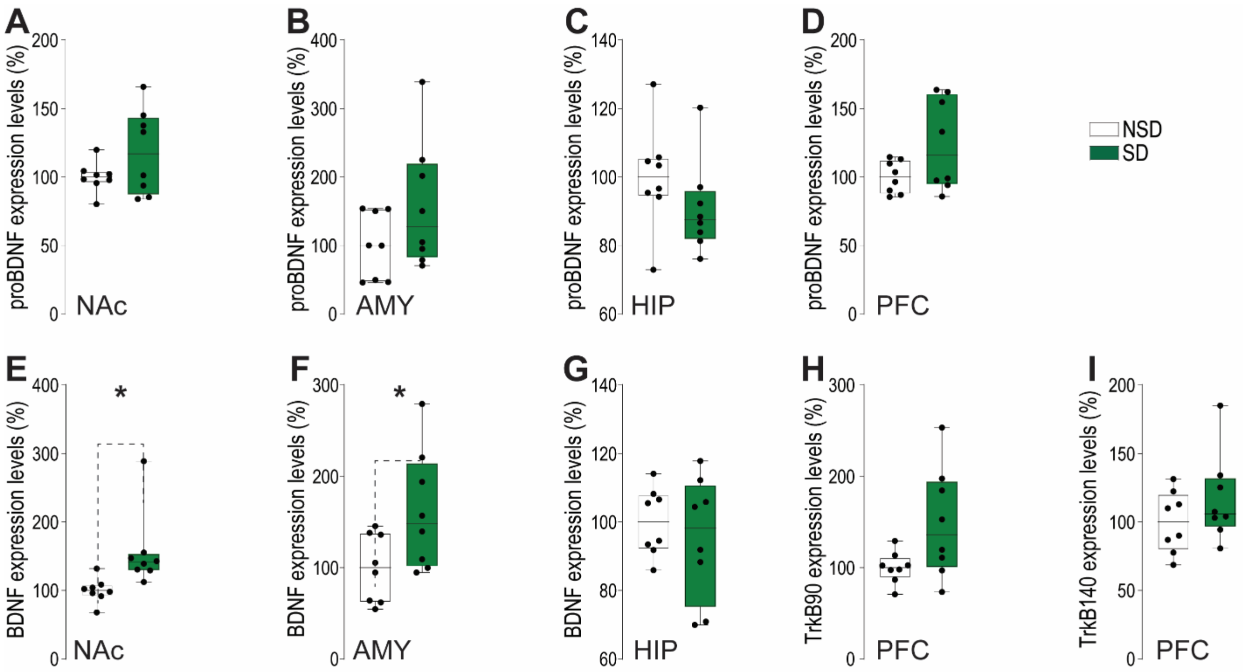


**Supplementary Figure 3 (Related to Figure 4).** Immediately following sleep deprivation (SD), rats were sacrificed and brain tissue was collected and flash-frozen. Western blot analyses of total lysates revealed that SD did not alter proBDNF expression in the nucleus accumbens (NAc, A), amygdala (AMY, B), hippocampus (HIP, C), or prefrontal cortex (PFC, D). In contrast, SD increased BDNF levels in the NAc (E) and AMY (F), but not in the HIP (G). Moreover, SD had no effect on TrkB receptor protein levels, either in the truncated (TrkB90; H) or full-length (TrkB140; I) forms. Data are presented as mean ± SEM and were analyzed using unpaired *t* tests with Welch's correction where appropriate. *, $p < 0.05$ for comparisons indicated by brackets. Each dot represents an individual animal.

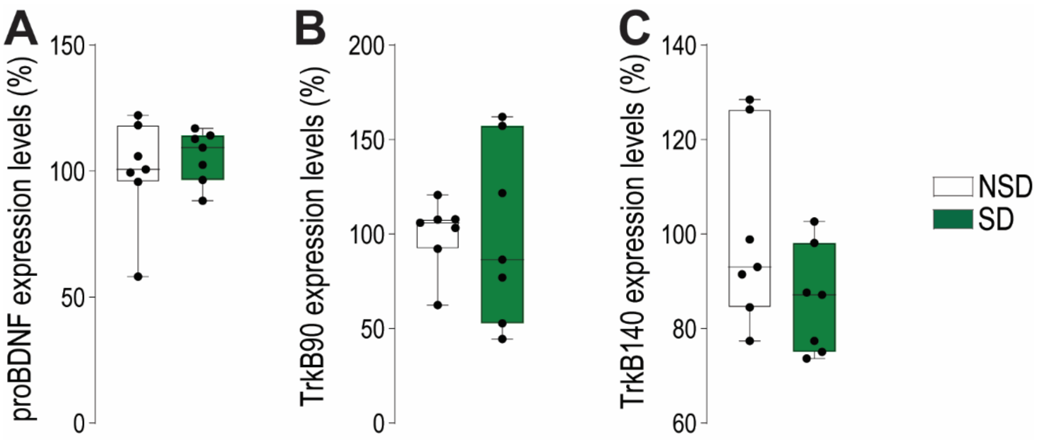


**Supplementary Figure 4 (Related to Figure 4).** Immediately following sleep deprivation (SD), mice were sacrificed and brain tissue was rapidly collected and flash-frozen. Western blot analyses of the prefrontal cortex showed that SD did not alter proBDNF expression (A) or TrkB receptor protein levels, either in the truncated (TrkB90; B) or full-length (TrkB140; C) forms. Data are presented as mean ± SEM and were analyzed using unpaired t tests with Welch's correction where appropriate. *, $p < 0.05$ for comparisons indicated by brackets. Each dot represents an individual animal.

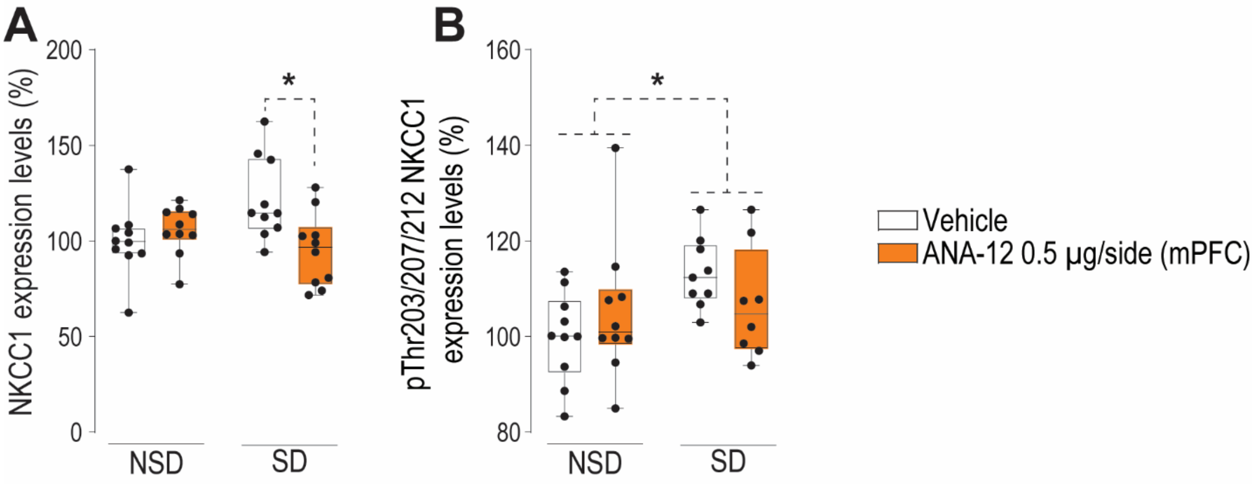


**Supplementary Figure 5 (Related to Figure 5).** Immediately following sleep deprivation (SD), rats were sacrificed and brain tissue was collected and flash-frozen. Western blot analyses of membrane-enriched fractions revealed that intra–medial prefrontal cortex (mPFC) administration of ANA-12 (0.5 µg/side) reduced NKCC1 membrane expression in SD rats (A), without altering the SD-induced decrease in NKCC1 phosphorylation (B), relative to non–sleep-deprived (NSD) controls. Data are presented as mean ± SEM and were analyzed using two-way ANOVA. *, $p < 0.05$ for comparison indicated by brackets. Each dot represents an individual animal.

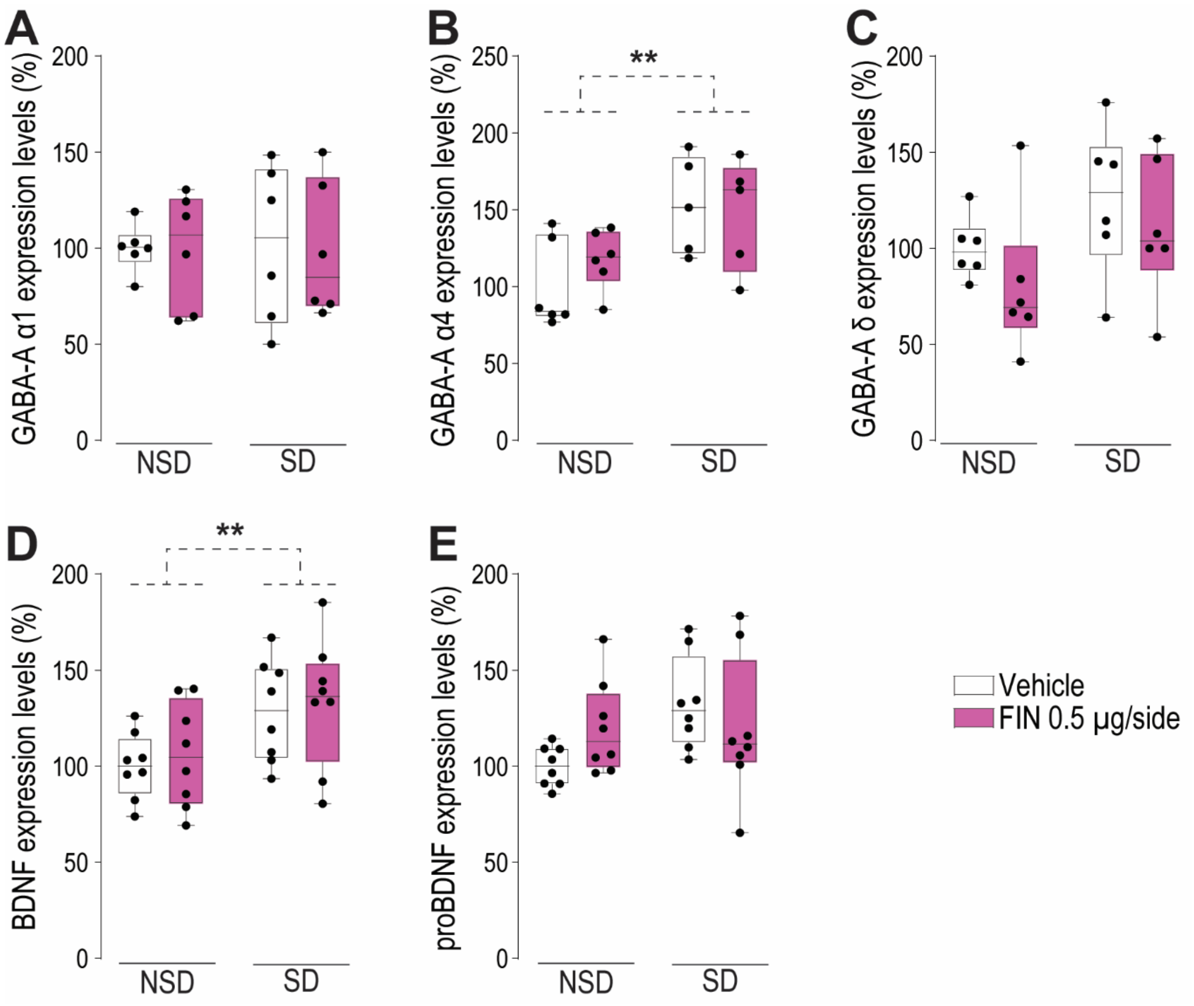


**Supplementary Figure 6 (Related to Figure 6).** Immediately following sleep deprivation (SD), rats were sacrificed and brain tissue was collected and flash-frozen. Western blot analyses showed that intra–medial prefrontal cortex (mPFC) administration of finasteride (FIN) did not affect GABA-A receptor subunits in membrane-enriched fractions (A–C). Specifically, no differences were observed in the α1 (A) or δ (C) subunits, whereas only SD-induced changes were detected in the α4 subunit (B). In total lysates, FIN (0.5 µg per side) did not modify SD-induced alterations in mature BDNF levels (D) and had no effect on proBDNF expression (E) relative to non–sleep-deprived (NSD) controls. Data are presented as mean ± s.e.m. and were analyzed using two-way ANOVA. **, $p < 0.01$ for the comparison indicated by brackets. Each dot represents one animal.

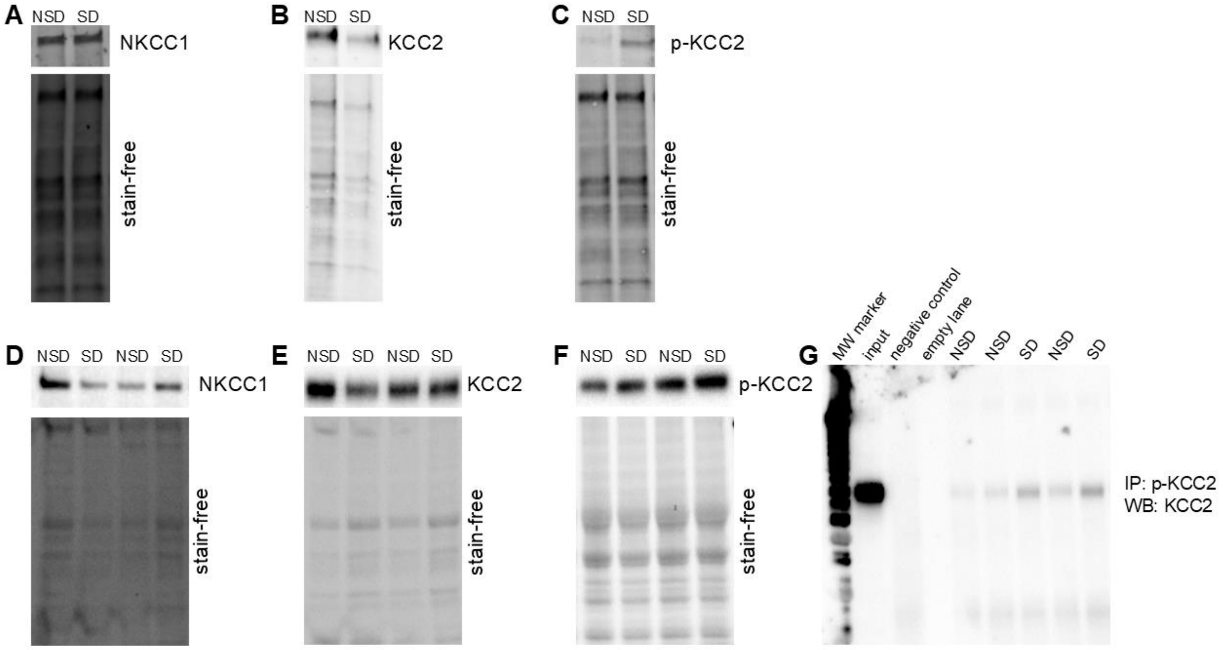


**Supplementary Figure 7 (Related to Figure 1).** Representative Western blot images used for quantification of NKCC1 (A, D), KCC2 (B, E), and KCC2 phosphorylated at Thr1007 (C, F) in membrane-enriched fractions from mice (A–C) and rats (D–F) under control (NSD) and sleep deprivation (SD) conditions. The corresponding stain-free images used for normalization are shown beneath each panel. Panel G shows immunoprecipitation performed using p-KCC2 (Thr1007) and detected with total KCC2.

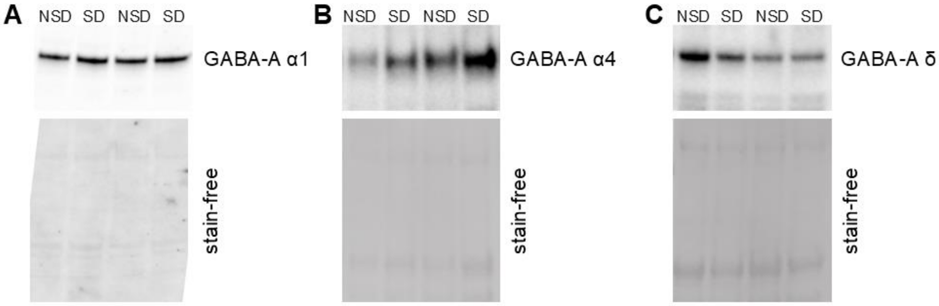


**Supplementary Figure 8 (Related to Figure 4).** Representative Western blot images used for quantification of the different GABA-A receptor subunits α1 (A), α4 (B), and δ (C) in membrane-enriched fractions from rats under control (NSD) and sleep deprivation (SD) conditions. Corresponding stain-free images used for normalization are shown beneath each panel.

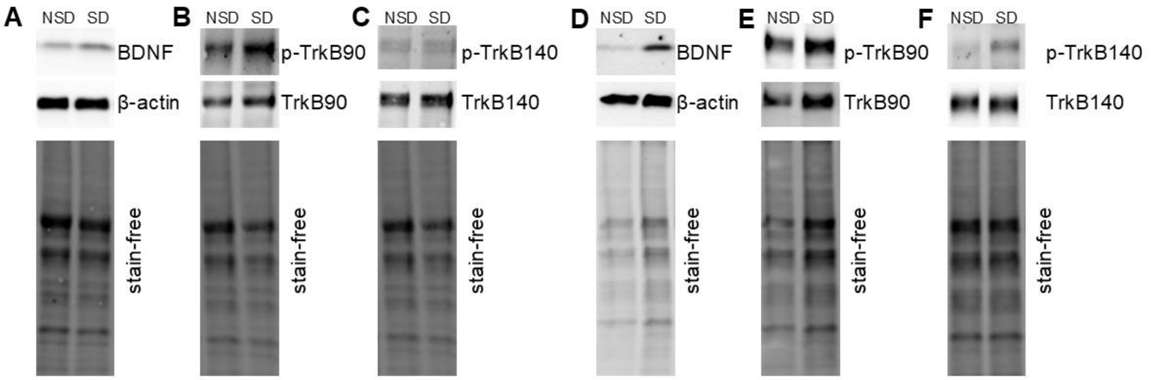


**Supplementary Figure 9 (Related to Figure 4).** Representative Western blot images used for quantification of BDNF (A, D), truncated TrkB (TrkB90; phosphorylated and total) (B, E), and full-length TrkB (TrkB140; phosphorylated and total) (C, F) in total lysates from rats (A–C) and mice (D–F) under control (NSD) and sleep deprivation (SD) conditions. Representative images of β-actin are also shown for panels A and D. Corresponding stain-free images used for normalization are shown beneath each panel.

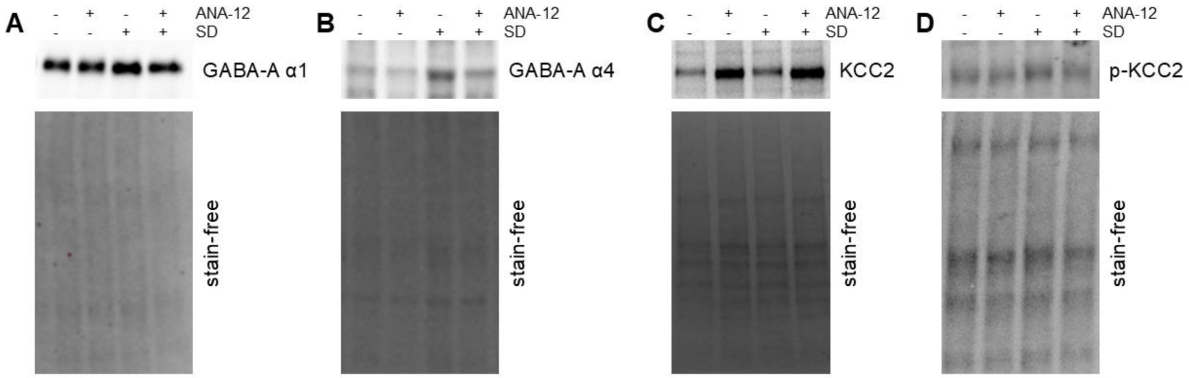


**Supplementary Figure 10 (Related to Figure 5).** Representative Western blot images used for quantification of the GABA-A receptor subunits α1 (A) and α4 (B), as well as total KCC2 (C) and phosphorylated KCC2 at Thr1007 (D), in membrane enriched fractions from rats under control and sleep deprivation (SD) conditions with or without treatment with the TrkB antagonist ANA-12. Corresponding stain-free images used for normalization are shown beneath each panel.

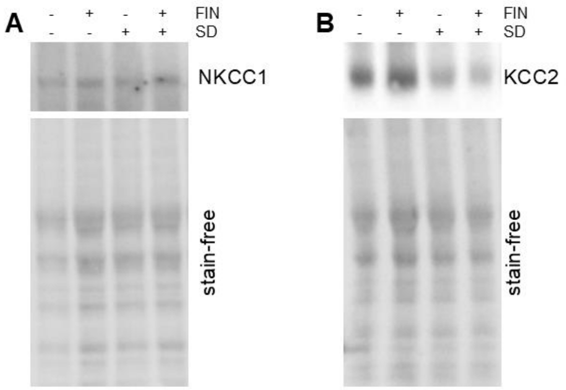


**Supplementary Figure 11 (Related to Figure 6).** Representative Western blot images used for quantification of NKCC1 (A) and KCC2 (B), in membrane-enriched fractions from rats under control and sleep deprivation (SD) conditions with or without treatment with finasteride (FIN). Corresponding stain-free images used for normalization are shown beneath each panel.

### SUPPLEMENTARY RESULTS

**SD Alters the Expression and Activity of Chloride Transporters NKCC1 and KCC2 in the PFC.** We explored whether the SD elicited changes in the expression or activity of NKCC1 and KCC2, whose functions are bidirectionally regulated by phosphorylation. Specifically, KCC2 activity is enhanced by phosphorylation at Ser940(Lee et al., 2007; Leonzino et al., 2016), whereas phosphorylation at Thr1007 reduces its activity by promoting internalization from the plasma membrane(Pracucci et al., 2023; Rinehart et al., 2009; Silayeva et al., 2015). In contrast, NKCC1 activity is upregulated by phosphorylation at Thr203/Thr207/Thr212(Kahle et al., 2013; Rinehart et al., 2009).

Based on this background, we examined the effects of SD on the expression and phosphorylation of the chloride transporters KCC2 and NKCC1 using biochemical fractionation and Western blot analysis. In mice, SD did not affect membrane-associated NKCC1 levels (**Fig. 1A**) [n = 7/group; t(12) = 0.10, p = 0.918, *d* = 0.06; two-tailed unpaired t test], whereas membrane KCC2 levels were significantly reduced following SD (**Fig. 1B**) [n = 7/group, t(12) = 2.71, p = 0.019, *d* = 1.45; two-tailed unpaired t test]. Consistent with this reduction, SD increased phosphorylation of membrane-associated KCC2 at Thr1007 (**Fig. 1C**) [n = 7/group, t(12) = 2.56, p = 0.025, *d* = 1.37; two-tailed unpaired t test], a modification known to impair membrane trafficking and reduce chloride extrusion capacity(Silayeva et al., 2015).

These effects were replicated in rats. As in mice, SD did not alter membrane-associated NKCC1 levels (**Fig. 1D**) [n = 8/group, t(14) = 0.48, p = 0.63 *d* = 0.24; unpaired t-test], while significantly decreasing membrane KCC2 levels (**Fig. 1E**) [n = 6/group, t(10) = 4.86, p = 0.0007, *d* = 2.80; unpaired t test]. Importantly, SD did not affect total protein abundance of either transporter, as neither total NKCC1 levels (**Fig. S1A**) [n = 5/group, t(8) = 0.18, p = 0.857, *d* = 0.12; unpaired t test] nor total KCC2 (**Fig. S1B**) [n = 5/group, t(8) = 1.55, p = 0.160, *d* = 0.98; unpaired t test] were significantly altered. These findings indicate that SD does not change chloride transporter abundance but instead modulates their function through post-translational mechanisms.

In line with this interpretation, phosphorylation analyses showed that SD selectively increased the phosphorylation state of NKCC1 at Thr203/207/212 (**Fig. S1C**) [n =

8/group, t(14) = 3.23, p = 0.006, *d* = 1.61; two-tailed unpaired t test], a modification known to enhance transporter activity and chloride import capacity. In addition, phosphorylation analyses further showed that SD selectively altered the phosphorylation state of KCC2: phosphorylation at Ser940 was unchanged (**Fig. S1D**) [n = 5/group, t(8) = 1.65, p = 0.137, *d* = 1.04; two-tailed unpaired t test], whereas phosphorylation at Thr1007 was significantly increased (**Fig. 1F**) [n = 8/group, t(14) = 2.28, p = 0.039, *d* = 1.14; two-tailed unpaired t test]. To directly confirm that SD-induced reductions in membrane KCC2 were mediated by phosphorylation at Thr1007, we performed immunoprecipitation of membrane fractions from mPFC lysates using a phospho-Thr1007-specific antibody, followed by immunoblotting for KCC2. SD significantly increased pThr1007-KCC2 levels in the membrane fraction (**Fig. 1G**) [n = 7-8/group, t(13) = 2.27, p = 0.040, *d* = 1.18], confirming that phosphorylated KCC2 is selectively removed from the membrane following SD. Taken together, these data indicate that SD disrupts chloride homeostasis through a dual mechanism: enhanced NKCC1 activity driven by increased phosphorylation promotes chloride import, while phosphorylation-dependent internalization of KCC2 reduces chloride extrusion. This combined effect leads to elevated intracellular chloride levels, depolarization of $E_{\_GABA}$, and compromised GABAergic inhibition in mPFC pyramidal neurons.

Given the established influence of circadian rhythms on cortical chloride homeostasis(Pracucci et al., 2023), we next examined whether SD-induced changes in NKCC1 and KCC2 regulation varied across diurnal and nocturnal phases. As shown in **Figure S2**, SD induced comparable alterations in chloride transporter expression and phosphorylation during both the light and dark phases. Specifically, SD produced significant main effects on membrane KCC2 expression (**Fig. S2B**) [n = 6/group; main effect of SD: F(1,20) = 7.53, p = 0.013, $\eta_p^2$ = 0.27; main effect of cycle: F(1,20) = 1.80, p = 0.194, $\eta_p^2$ = 0.08; interaction: F(1,20) = 0.004, p = 0.950, $\eta_p^2$ = 2.04x$10^{-4}$; two-way ANOVA], NKCC1 phosphorylation (**Fig. S2C**) [n = 7-8/group; main effect of SD: F(1,27) = 8.94, p = 0.006, $\eta_p^2$ = 0.25; main effect of cycle: F(1,27) = 1.81, p = 0.190, $\eta_p^2$ = 0.06; interaction: F(1,27) = 1.75, p = 0.197, $\eta_p^2$ = 0.06; two-way ANOVA], and KCC2 phosphorylation (**Fig. S2D**) [n = 5-8/group; main effect of SD: F(1,23) = 6.12, p = 0.021,

$\eta_p^2$ = 0.21; main effect of cycle: F(1,23) = 6.36, p = 0.019, $\eta_p^2$ = 0.22; interaction: F(1,23) = 0.11, p = 0.740, $\eta_p^2$ = 0.005; two-way ANOVA]. In contrast, no significant effects were detected for membrane NKCC1 expression (**Fig, S2A**) [n = 6-8/group; main effect of SD: F(1,25) = 0.14, p = 0.709, $\eta_p^2$ = 0.006; main effect of cycle: F(1,25) = 0.68, p = 0.416, $\eta_p^2$ = 0.03; interaction: F(1,25) = 0.07, p = 0.798, $\eta_p^2$ = 0.003; two-way ANOVA]. No significant interactions between SD and circadian phase were detected, indicating that SD-induced dysregulation of chloride transporter phosphorylation and membrane localization occurs independently of the circadian cycle.

**Sleep Deprivation Increases Intracellular Chloride Levels and Excitability of Cortical Pyramidal Neurons.** To investigate the impact of SD on intracellular chloride ($[Cl^-]_i$) homeostasis in cortical pyramidal neurons, we employed the pH- and chloride-sensitive biosensor LSSmClopHensor(Arosio et al., 2010). This construct was delivered via a floxed adeno-associated viral (AAV) vector into mice expressing Cre-recombinase under the Emx1 promoter, which drives expression selectively in excitatory cortical neurons(Sulis Sato et al., 2017). Previous studies have demonstrated that this approach enables selective labeling of supragranular (layer 2/3) excitatory pyramidal neurons in the cortex of young mice (1–4 months old), facilitating in vivo imaging through a cranial window with two-photon microscopy(Pracucci et al., 2023).

Using this approach, intracellular chloride concentrations ($[Cl^-]_i$) were measured in pyramidal neurons in cortical layers 2/3 under normal sleep (NSD) and SD conditions by two-photon imaging (**Fig. 2A**). While intracellular pH remained stable across conditions (NSD, pH = 7.20 *vs* SD, pH = 7.29, Mann-Whitney test), we observed a significant increase in $[Cl^-]_i$ in pyramidal neurons of SD-subjected mice compared to NSD controls (**Fig. 2B-C**, U = 3.0, p = 0.008, $r_{(rb)}$ = 0.83, Mann-Whitney test).

Since GABA-A receptors are primarily permeable to chloride, their equilibrium potential ($E_{\_GABA}$) is dictated by transmembrane chloride gradients. More negative $E_{\_GABA}$ values enhance synaptic inhibition via hyperpolarization, whereas more positive $E_{\_GABA}$ values diminish inhibition and may facilitate depolarization, potentially shifting GABAergic signaling from inhibitory to excitatory. To assess whether increased $[Cl^-]_i$ in mPFC pyramidal neurons of SD rats corresponded to a depolarizing shift in $E_{\_GABA}$, we utilized the gramicidin-perforated patch-clamp technique, which preserves native intracellular chloride concentrations while enabling measurement of GABA-A-mediated currents at varying membrane potentials(Kyrozis & Reichling, 1995). Consistent with prior findings(Alfonsa et al., 2023), current-voltage (I-V) curves for GABA-A-mediated currents significantly differed between SD and NSD rats (**Fig. 2D**) [interaction: SD condition × voltage: $F(7, 98) = 2.4$, $p = 0.026$; main effect of SD: $F(2.561, 35.85) = 7.88$, $p = 0.0006$, two-way repeated-measure ANOVA]. Importantly, SD induced a significant depolarizing shift in $E_{\_GABA}$ toward more positive values (**Fig. 2E**) [$t(22.49) = 2.91$, $p = 0.008$, $d = 1.04$, unpaired t-test with Welch's correction], consistent with elevated $[Cl^-]_i$.

We next examined whether SD-mediated depolarizing shifts in $E_{\_GABA}$ affected the intrinsic excitability of pyramidal neurons within the prelimbic mPFC (layer II-III) using whole-cell patch-clamp electrophysiology. Current-clamp recordings revealed that SD pyramidal neurons exhibited significantly increased firing frequency in response to somatically injected depolarizing currents (**Fig. 2F**) [main effect of SD: $F(1, 12) = 6.67$, $p = 0.024$, $\eta_p^2 = 0.36$; main effect of stimulus intensity: $F(1.446, 17.36) = 42.70$, $p < 0.0001$, $\eta_p^2 = 0.78$; interaction: $F(1.446, 17.36) = 6.97$, $p = 0.01$, $\eta_p^2 = 0.37$, two-way repeated measures ANOVA with Greenhouse-Geisser correction]. Additionally, SD neurons displayed a higher maximal firing rate (**Fig. 2G**) [$t\ (8.146) = 3.26$, $p = 0.011$, $d = 1.65$, unpaired t-test with Welch's correction], a lower rheobase (the minimal current intensity required to evoke an action potential) (**Fig. 2H**) [$U = 39$, $p = 0.009$, $r_{(rb)} = 0.86$, non-parametric Mann–Whitney U test], and a decreased voltage threshold for action potential initiation (**Fig. 2I**) [$U = 40$, $p = 0.043$, $r_{(rb)} = 0.67$, non-parametric Mann–Whitney U test]. These findings collectively indicate enhanced intrinsic excitability of mPFC pyramidal neurons following SD.

To evaluate the effects of SD on synaptic inhibition, we measured the paired-pulse ratio (PPR) of GABA-A-mediated inhibitory postsynaptic currents (IPSCs) in mPFC pyramidal neurons. The PPR is an indicator of presynaptic release probability and short-term synaptic plasticity, with reduced PPR suggesting enhanced neurotransmitter release or altered postsynaptic receptor function. SD significantly reduced the PPR compared to

NSD (**Fig. 2J**) [t(12) = 2.59, p = 0.024, *d* = 1.38, unpaired t-test], indicating altered GABAergic synaptic transmission.

Taken together, these findings demonstrate that SD disrupts chloride homeostasis and increases neuronal excitability in the mPFC, leading to alterations in both intrinsic and synaptic properties of pyramidal neurons. These effects were consistently observed in both rats and mice, suggesting a conserved mechanism by which SD compromises cortical inhibitory function.

**The NKCC1 Inhibitor Bumetanide Rescues Neurobehavioral Complications Induced by SD.** We previously demonstrated that, in both rats and mice, SD elicits a broad spectrum of cognitive and behavioral complications under the control of the mPFC, including deficits in executive functions, sensorimotor gating, and impulse control (Cadeddu et al., 2022; Frau et al., 2008, 2017). Since the pathological rise in $[Cl^-]_i$ within PFC pyramidal neurons can be mitigated by bumetanide, an inhibitor of NKCC1 (Pracucci et al., 2023; Sulis Sato et al., 2017), we investigated the effect of this drug on the impairment of information processing elicited by SD in mice and rats *in vivo* (25-50 mg/kg).

In male mice, SD did not alter the acoustic startle reflex (**Fig. 3A**) [n = 8/group; main effect of SD: F(1,42) = 0.76, p = 0.387, $\eta_p^2$ = 0.02]. In parallel, we confirmed that SD produces robust deficits in PPI (**Fig. 3B**) [main effect of SD: F(1,42) = 20.62, p < 0.0001, $\eta_p^2$ = 0.33], a reliable operational measure of sensorimotor gating that is impaired in various neuropsychiatric disorders (Frau et al., 2008, 2017). Bumetanide systemic administration did not affect startle amplitude in either NSD or SD male mice [main effect of treatment: F(2,42) = 0.74, p = 0.483, $\eta_p^2$ = 0.03; interaction: F(2,42) = 0.88, p = 0.421, $\eta_p^2$ = 0.04].

Notably, systemic administration of bumetanide fully reversed the SD-induced PPI impairments [main effect of treatment: F(2,42) = 9.31, p = 0.0004, $\eta_p^2$ = 0.31; interaction: F(2,42) = 12.98, p < 0.0001, $\eta_p^2$ = 0.38].

Similar results were obtained in female mice, where neither SD nor bumetanide treatment altered the startle amplitude (**Fig. 3C**) [n= 8/group; main effect of SD: F(1,42) = 0.09, p = 0.763, $\eta_p^2$ = 0.002; main effect of treatment: F(2,42) = 0.19, p = 0.826, $\eta_p^2$ = 0.009;

interaction: F(2,42) = 0.07, p = 0.933, $\eta_p^2$ = 0.003], while systemic administration of bumetanide at the higher dose (50 mg/kg) fully reversed the SD-induced PPI impairments (**Fig. 3D**) [main effect of SD: F(1,42) = 6.06, p = 0.018, $\eta_p^2$ = 0.13; main effect of treatment: F(2,42) = 2.98, p = 0.062, $\eta_p^2$ = 0.12; interaction: F(2,42) = 3.26, p = 0.048, $\eta_p^2$ = 0.13].

Similar results were obtained in rats. SD, irrespective of bumetanide treatment, significantly altered the acoustic startle reflex (**Fig. 3E**) [n = 8-17/group; main effect of SD: F(1,57) = 11.82, p = 0.001, $\eta_p^2$ = 0.17; main effect of treatment: F(2,57) = 1.61, p = 0.208, $\eta_p^2$ = 0.05; interaction: F(2,57) = 0.39, p = 0.680, $\eta_p^2$ = 0.01]. In parallel, we confirmed that SD produces robust deficits in PPI (**Fig. 3F**) [main effect of SD: F(1,57) = 14.95, p = 0.0003, $\eta_p^2$ = 0.21]. Notably, systemic administration of bumetanide dose-dependently reversed the SD-induced PPI impairments (**Fig. 3G**) [main effect of treatment: F(2,57) = 1.78, p = 0.179, $\eta_p^2$ = 0.06; interaction: F(2,57) = 3.91, p = 0.026, , $\eta_p^2$ = 0.12], with the higher dose (50 mg/kg) fully normalizing PPI to NSD levels.

In the novel object recognition (NOR) test, which assesses non-spatial declarative memory dependent on mPFC function (Barker & Warburton, 2011), SD caused significant long-term deficits in memory recognition of the novel object, as indicated by a marked reduction in the novel exploration index (%NEI) (**Fig. 3H**) [n = 8/group; main effect of SD: F(1,28) = 39.40, p < 0.0001, $\eta_p^2$ = 0.58]. Consistent with the PPI findings, bumetanide systemic administration effectively restored the %NEI in SD rats [main effect of treatment: F(1,28) = 12.52, p = 0.001, $\eta_p^2$ = 0.31; interaction: F(1,28) = 8.84, p = 0.006, $\eta_p^2$ = 0.24], indicating complete rescue of recognition memory deficits.

These findings indicate that pharmacological inhibition of NKCC1 with bumetanide is sufficient to prevent SD-induced cognitive and behavioral deficits, supporting a causal role for elevated $[Cl^-]_i$ and disrupted chloride homeostasis in the pathophysiology of SD-related information processing impairments.

**SD Alters GABA-A Receptor Subunit Composition in the mPFC.** We next investigated whether SD may also alter GABA-A receptor expression and subunit composition s in the mPFC (**Fig. 4A-C**). Western blot analysis revealed that SD-subjected rats exhibited a selective and significant increase in the GABA-A α4 subunit (**Fig. 4B**) [n = 6/group, t(10)

= 2.69, p = 0.023, *d* = 1.55, unpaired t-test], while no significant changes were observed in the levels of GABA-A α1 (**Fig. 4A**) [n = 5-7/group, t(6.793) = 0.54, p = 0.605, *d* = 0.29, unpaired t-test with Welch's correction], GABA-A δ (**Fig. 4C**) [n = 5-7/group, t(10) = 0.88, p = 0.399, *d* = 0.52, unpaired t-test]. The α4 subunit-containing GABA-A receptors are typically extrasynaptic, exhibit high affinity for GABA, and are particularly sensitive to neurosteroid modulation (Belelli et al., 2009). Moreover, α4-containing receptors can mediate tonic inhibition and are associated with altered neuronal excitability in various pathological states (Maguire & Mody, 2008). The selective upregulation of GABA-A α4 following SD may represent a compensatory response to elevated neurosteroid levels and disrupted chloride homeostasis, potentially contributing to the altered inhibitory tone observed in mPFC pyramidal neurons.

We next investigated the upstream mechanisms potentially linking the observed increase in GABA-A receptor α4 subunit expression with the reduction in membrane-associated KCC2. Among the candidates, brain-derived neurotrophic factor (BDNF) was of particular interest, given its well-established involvement in regulating GABA-A receptor subunit composition and KCC2 membrane trafficking, as well as prior reports demonstrating elevated BDNF levels following sleep deprivation. No changes in proBDNF levels were detected in any of the regions examined (**Fig. S3A-D**) [n = 8/group; nucleus accumbens (NAc): t(8.72) = 1.56, p = 0.153, *d* = 0.78; amygdala (AMY): t(14) = 1.57, p = 0.138, *d* = 0.79; hippocampus (HIP): t(14) = 1.29, p = 0.218, *d* = 0.64; PFC: t(8.74) = 1.90, p = 0.091, *d* = 0.95; unpaired t-test, with Welch's correction applied for PFC and NAc]. Notably, SD in rats produced a robust increase in mature BDNF levels across multiple brain regions, including PFC (**Fig. 4D**) [n = 8/group, t(14) = 2.45, p = 0.028, *d* = 1.22, unpaired t-test], NAc (**Fig. S3E**) [n = 8/group, t(8.44) = 2.71, p = 0.025, *d* = 1.36, unpaired t-test with Welch's correction], and AMY (**Fig. S3F**) [n = 8/group, t(14) = 2.32, p = 0.036, *d* = 1.16, unpaired t-test], whereas no changes were observed in the HIP (**Fig. S3C**) [t(14) = 0.67, p = 0.516, *d* = 0.33, unpaired t-test]. In addition, rats exposed to SD did not show any alterations in the phosphorylation state of TrkB, either in its truncated (**Fig. 4E**) [n = 8/group, t(14) = 0.64, p = 0.533, *d* = 0.32, unpaired t-test] or full-length form (**Fig. 4F**) [n = 8/group, t(14) = 1.53, p = 0.147, *d* = 0.77, unpaired t-test]. Similarly, SD did not significantly affect total TrkB protein levels, although a trend toward increased expression of the truncated TrkB isoform was observed (**Fig. S3H**) [n = 8/group, t(8.15) = 2.20, p = 0.059, *d* = 1.10; unpaired t test with Welch's correction), while full-length TrkB levels remained unchanged (**Fig. S3I**) [n = 8/group, t(14) = 1.21, p = 0.247, *d* = 0.60, unpaired t test).

Parallel results were obtained in mice, where SD induced a robust increase in mature BDNF in PFC (**Fig. 4G-H**) [panel H: n = 7/group, t(12) = 2.74, p = 0.018, *d* = 1.46, unpaired t test] without affecting proBDNF expression (**Fig. S4A**) [n = 7/group, t(12) = 0.65, p=0.529, *d* = 0.35, unpaired t test]. Analysis of TrkB signaling revealed a selective increase in phosphorylation of the full length receptor, with no change in the truncated form (**Fig. 4I-J**) [n = 7/group; TrkB90: t(6.30) = 1.88, p = 0.107, *d* = 1.00; TrkB140: t(7.90) = 3.23, p = 0.012, *d* = 1.73, unpaired t test with Welch's correction]. Notably, SD did not alter total TrkB protein levels in either isoform (**Fig. S4B-C**) [n = 7/group; TrkB90: t(7.78) = 0.01, p = 0.990, *d* = 0.01; TrkB140: t(12) = 1.62, p = 0.131, *d* = 0.87, unpaired t test with Welch's correction applied to TrkB90].

**TrkB blockade mitigates SD-induced molecular and behavioral alterations.** To determine whether enhanced BDNF–TrkB signaling contributes causally to the information-processing deficits elicited by SD, we inhibited TrkB signaling using the selective antagonist ANA-12, administered either systemically (1 mg/kg, **Fig. 5A–B**) or locally into the mPFC (0.5µg/0.5µl/side, **Fig. 5C–J**). Neither SD or systemic administration of ANA-12 alter baseline startle amplitude (**Fig. 5A**) [n = 8/group; main effect of treatment: F(1,32) = 0.14, p = 0.713, $\eta_p^2$ = 0.004; main effect of SD: F(1,32) = 0.21, p = 0.650, $\eta_p^2$ = 0.006; interaction: F(1,32) = 0.0002, p = 0.990, $\eta_p^2$ = 4.79x10$^{-6}$; two-way ANOVA]. In contrast, systemic ANA-12 produced a partial, near-significant attenuation of SD-induced PPI deficits (**Fig. 5B**) [main effect of treatment: F(1,32) = 7.95, p = 0.008, $\eta_p^2$ = 0.20; main effect of SD: F(1,32) = 13.54, p = 0.0008, $\eta_p^2$ = 0.30; interaction: F(1,32) = 2.86, p = 0.100, $\eta_p^2$ = 0.08; two-way ANOVA]. In contrast, while intra–mPFC delivery of ANA-12 did not alter mean startle amplitude (**Fig. 5C**) [n = 12-13/group; main effect of treatment: F(1,47) = 0.96, p = 0.331, $\eta_p^2$ = 0.02; main effect of SD F(1,47) = 1.24, p = 0.271, $\eta_p^2$ = 0.03; interaction F(1,47) = 0.26, p = 0.611, $\eta_p^2$ = 0.006; two-way ANOVA], it fully prevented SD-induced PPI deficits (**Fig. 5D**), evidenced by a significant treatment

× SD interaction [F(1,47) = 5.13, p = 0.028, $\eta_p^2$ = 0.10], alongside a main effect of SD [F(1,47) = 12.95, p = 0.0008, $\eta_p^2$ = 0.22] and no significant main effect of treatment [F(1,47) = 2.94, p = 0.093, $\eta_p^2$ = 0.059; two-way ANOVA]. Moreover, intra-mPFC administration of ANA-12 effectively restored %NEI (**Fig. 5E**) in SD rats, as indicated by significant main effects of treatment [n = 10-14/group; F(1,45) = 9.53, p = 0.004, $\eta_p^2$ = 0.18] and SD condition [F(1,45) = 13.98, p = 0.0005, $\eta_p^2$ = 0.24], as well as a significant treatment × SD interaction [F(1,45) = 6.27, p = 0.016, $\eta_p^2$ = 0.12; two-way ANOVA], indicating a complete rescue of SD-induced recognition memory deficits.

At the molecular level, ANA-12 administration did not affect GABA-A α1 levels (**Fig. 5F**) [n=6/group; main effect of treatment: F(1,20) = 1.77, p = 0.198, $\eta_p^2$ = 0.08; main effect of SD: F(1,20) = 3.56, p = 0.074, $\eta_p^2$ = 0.15; interaction: F(1,20) = 2.63, p = 0.120, $\eta_p^2$ = 0.12; two-way ANOVA]. In contrast, ANA-12 effectively normalized the SD-induced upregulation of the GABA-A receptor α4 subunit (**Fig. 5G**), as evidenced by a significant treatment × SD interaction [n= 6-7/group; F(1,22) = 14.12, *p* = 0.001, $\eta_p^2$ = 0.39], in the absence of significant main effects of treatment or SD condition [treatment: F(1,22) = 0.41, *p* = 0.527, $\eta_p^2$ = 0.02; SD: F(1,22) = 3.22, *p* = 0.086, $\eta_p^2$ = 0.13; two-way ANOVA]. The behavioral rescue was accompanied by coordinated changes in chloride transporters. Intra-mPFC administration of ANA-12 reduced NKKC1 membrane expression in SD rats (**Fig. S5A**) [n = 10/group; main effect of treatment: F(1,36) = 3.21, p = 0.081, $\eta_p^2$ = 0.08; main effect of SD: F(1,36) = 0.92, p = 0.344, $\eta_p^2$ = 0.02; interaction: F(1,36) = 7.71, p = 0.009, $\eta_p^2$ = 0.18; two-way ANOVA], without altering the SD-induced reduction in NKCC1 phosphorylation (**Fig. S5B**) [n = 8-10/group; main effect of treatment: F(1,33) = 0.03, p = 0.866, $\eta_p^2$ = 8.70x10$^{-4}$; main effect of SD: F(1,33) = 4.15, p = 0.050, $\eta_p^2$ = 0.11; interaction: F(1,33) = 2.38, p = 0.132, $\eta_p^2$ = 0.07; two-way ANOVA]. In parallel, ANA-12 restored KCC2 membrane expression (**Fig. 5H**) [n = 8/group; main effect of treatment: F(1,28) = 4.72, p = 0.038, $\eta_p^2$ = 0.14; main effect of SD: F(1,28) = 2.59, p = 0.119, $\eta_p^2$ = 0.08; interaction: F(1,28) = 5.75, p = 0.023, $\eta_p^2$ = 0.17; two-way ANOVA] and normalized SD-induced hyperphosphorylation (**Fig. 5I**) [n = 8/group; main effect of treatment: F(1,28) = 5.37, p = 0.028, $\eta_p^2$ = 0.16; main effect of SD: F(1,28) = 1.98, p = 0.170, $\eta_p^2$ = 0.07; interaction: F(1,28) = 6.61, p = 0.016, $\eta_p^2$ = 0.19; two-way ANOVA].

**The 5α-Reductase Inhibitor Finasteride Rescues Neurobehavioral Complications Induced by SD.** We previously reported that the behavioral (endo)phenotypes induced by SD are mediated by elevated AP levels in the PFC and are further exacerbated by exogenous AP administration (Frau et al., 2017). Furthermore, we observed that PPI deficits inversely correlate with 5α-reductase (5αR) expression in the PFC, with higher enzyme levels associated with more pronounced PPI impairments (Frau et al., 2017). Building on these findings, we investigated whether irreversible blockade of 5αR using finasteride (FIN), which depletes AP levels, could restore neurosteroid homeostasis and ameliorate SD-induced deficits. FIN was administered systemically in mice (12.5-25 mg/kg, IP) and directly into the mPFC in rats (0.5 μg/0.5 μl/side).

In female mice, neither SD nor FIN treatment altered the acoustic startle reflex (**Fig. 6A**) [n = 8/group; main effect of SD: F(1,42) = 2.74, p = 0.106, $\eta_p^2$ = 0.06; main effect of treatment: F(2,42) = 1.01, p = 0.373, $\eta_p^2$ = 0.05; interaction: F(2,42) = 0.13, p = 0.882, $\eta_p^2$ = 0.006; two-way ANOVA]. Consistent with previous findings, SD induced robust deficits in PPI (**Fig. 6B**) [main effect of SD: F(1,42) = 14.73, p=0.0004, $\eta_p^2$ = 0.26]. Notably, FIN attenuated SD-induced PPI impairments, as reflected by a main effect of treatment [F(2,42) = 3.58, *p* = 0.037, $\eta_p^2$ = 0.15] and a trend toward a treatment × SD interaction [F(2,42) = 2.67, *p* = 0.081, $\eta_p^2$ = 0.11; two-way ANOVA].

Similar effects were observed in male mice. Neither SD nor FIN treatment affected the startle amplitude (**Fig. 6C**) [n = 8/group; main effect of SD: F(1,42) = 0.94, p = 0.339, $\eta_p^2$ = 0.02; main effect of treatment: F(2,42) = 0.99, p = 0.381, $\eta_p^2$ = 0.04; interaction: F(2,42) = 0.52, p = 0.600, $\eta_p^2$ = 0.02; two-way ANOVA]. In contrast, systemic FIN administration at the higher dose (25 mg/kg) fully reversed the SD-induced PPI impairments (**Fig. 6D**) [n = 8/group, main effect of SD: F(1,42) = 6.59, p = 0.014, , $\eta_p^2$ = 0.14; main effect of treatment: F(2,42) = 5.77, p = 0.006, $\eta_p^2$ = 0.22; interaction: F(2,42) = 5.37, p = 0.008, $\eta_p^2$ = 0.20; two-way ANOVA]. Importantly, these behavioral improvements were accompanied by increased in $[Cl^-]_i$, as assessed by two-photon imaging of LSSmClopHensor (**Fig. 6E,** $p < 0.05$, $r_{(rb)} = 0.8$, Mann-Whitney test).

In rats, neither SD nor intra–PFC infusion of FIN affected startle amplitude (**Fig. 6F**) [n = 10-11/group; main effect of SD: $F(1,37) = 3.64$, $p = 0.064$, $\eta_p^2 = 0.09$; main effect of FIN: $F(1,37) = 0.02$, $p = 0.879$, $\eta_p^2 = 6.34x10^{-4}$; interaction: $F(1,37) = 0.15$, $p = 0.703$, $\eta_p^2 = 0.004$; two-way ANOVA]. As expected, SD significantly impaired PPI (**Fig. 6G**) [main effect of SD: $F(1,37) = 15.60$, $p = 0.0003$, $\eta_p^2 = 0.30$], and this deficit was rescued by intra-PFC FIN administration, as indicated by a significant main effect of treatment [$F(1,37) = 4.52$, $p = 0.040$, $\eta_p^2 = 0.11$] and a trend toward a treatment × SD interaction [$F(1,37) = 3.33$, $p = 0.076$, $\eta_p^2 = 0.08$; two-way ANOVA]. Finally, SD induced robust deficits in novel object recognition memory, reflected by a reduced %NEI (**Fig. 6H**) [n = 9-10/group; main effect of SD: $F(1,35) = 25.36$, $p < 0.0001$, $\eta_p^2 = 0.42$]. FIN had no effect on %NEI in NSD rats [main effect of FIN: $F(1,35) = 1.48$, $p = 0.232$, $\eta_p^2 = 0.04$], but fully normalized the %NEI in SD rats [interaction: $F(1,35) = 28.58$, $p < 0.0001$, $\eta_p^2 = 0.45$; two-way ANOVA].

At the molecular level, neither SD nor FIN treatment altered NKCC1 membrane expression (**Fig. 6I**) [n = 6/group, main effect of SD: $F(1,20) = 0.65$, $p = 0.428$, $\eta_p^2 = 0.03$; main effect of treatment: $F(1,20) = 0.31$, $p = 0.584$, $\eta_p^2 = 0.02$; interaction: $F(1,20) = 1.28$, $p = 0.271$, $\eta_p^2 = 0.06$; two-way ANOVA]. In parallel, FIN intra-PFC administration did not modify the SD-induced reduction in KCC2 membrane expression (**Fig. 6J**) [n = 6/group; main effect of SD: $F(1,20) = 5.76$, $p = 0.026$, $\eta_p^2 = 0.22$; main effect of treatment: $F(1,20) = 0.008$, $p = 0.929$, $\eta_p^2 = 4.10x10^{-4}$; interaction: $F(1,20) = 1.18$, $p = 0.290$, , $\eta_p^2 = 0.06$; two-way ANOVA]. Consistent with these findings, further analyses of GABA-A subunits showed that FIN treatment did not affect relative subunits expression (**Fig. S6A-C**). Specifically, FIN had no effect on α1 subunit levels (**Fig. S6A**) [n = 6/group, main effect of SD: $F(1,20) = 0.002$, $p = 0.963$, $\eta_p^2 = 1.12x10^{-4}$; main effect of treatment: $F(1,20) = 0.03$, $p = 0.858$, $\eta_p^2 = 0.002$; interaction: $F(1,20) = 0.01$, $p = 0.909$, $\eta_p^2 = 6.76x10^{-4}$; two-way ANOVA] or on δ subunit expressions (**Fig. S6C**) [n= 6/group; main effect of SD: $F(1,20) = 4.02$, $p = 0.059$, $\eta_p^2 = 0.17$; main effect of treatment: $F(1,20) = 1.50$, $p = 0.235$, $\eta_p^2 = 0.07$; interaction: $F(1,20) = 0.04$, $p = 0.842$, $\eta_p^2 = 0.002$; two-way ANOVA]. Moreover, FIN failed to reverse the SD-induced upregulation of the α4 subunit (**Fig. S6B**) [n = 5-6/group; main effect of SD: F(1,18) = 10.82, p = 0.004, $\eta_p^2$ = 0.38; main effect of treatment: F(1,18) = 0.24, p = 0.630, $\eta_p^2$ = 0.01; interaction: F(1,18) = 0.86, p = 0.364, $\eta_p^2$ = 0.05; two-way ANOVA]. In addition, FIN treatment did not modify SD-induced changes in mature BDNF protein levels (**Fig. S6D**) [n = 8/group; main effect of SD: F(1,28) = 8.72, p = 0.006, $\eta_p^2$ = 0.24; main effect of treatment: F(1,28) = 0.29, p = 0.595, $\eta_p^2$ = 0.01; interaction: F(1,28) = 0.005, p = 0.944, $\eta_p^2$ = 1.81x10$^{-4}$; two-way ANOVA]. FIN also had no effect on proBDNF expression (**Fig. S6E**) [n = 8/group; main effect of SD: F(1,28) = 3.22, p = 0.084, $\eta_p^2$ = 0.10; main effect of treatment: F(1,28) = 0.14, p = 0.714, $\eta_p^2$ = 0.005; interaction: F(1,28) = 3.28, p = 0.081, $\eta_p^2$ = 0.10; two-way ANOVA]. These results suggest that the therapeutic effects of FIN are not mediated by alterations in the membrane distribution of chloride-cation cotransporters or changes in GABA-A receptor subunit composition. Instead, the rescue effects are likely attributable to AP depletion and the subsequent reduction in neurosteroid-mediated potentiation of depolarizing GABA-A receptor currents. This interpretation is consistent with the finding that SD-induced alterations in chloride transporter expression and GABA-A α4 upregulation persist despite FIN treatment, yet behavioral deficits are normalized, indicating that elevated AP levels are necessary for the functional consequences of disrupted chloride homeostasis to manifest as cognitive impairments.

**AP is Necessary but Not Sufficient to elicit SD-Like behavioral Deficits.** Finally, we examined whether AP was necessary and/or sufficient to induce an SD-like phenotype in NSD rats. To test this hypothesis, AP was administered either systemically (10 mg/kg, **Fig. 7A-B**) or directly into the mPFC (0.5 μg/0.5 μl/side, **Fig. 7C-D**), alone or in combination with the selective KCC2 inhibitor VU0463271 VU0463271 vu(VU; 0.02 μg/0.5 μl/side), which impairs chloride extrusion and elevates $[Cl^-]_i$ (Delpire et al., 2009; Sivakumaran et al., 2015).

Neither AP nor VU alone, nor their combination, altered baseline acoustic startle amplitude, irrespective of the route of AP delivery (**Fig. 7A,C**) [systemic AP experiment: n = 8/group; main effect of VU: F(1,28) = 0.40, p = 0.530, $\eta_p^2$ = 0.01; main effect of AP:

F(1,28) = 0.38, p = 0.540, $\eta_p^2$ = 0.01; interaction: F(1,28)=0.12, p = 0.735, $\eta_p^2$ = 0.004; mPFC AP experiment: n=8-11/group; main effect of VU: F(1,31) = 0.21, p = 0.648, $\eta_p^2$ = 0.007; main effect of AP: F(1,31) = 2.81, p = 0.104, $\eta_p^2$ = 0.08; interaction: F(1,31) = 0.93, p = 0.341, $\eta_p^2$ = 0.03; two-way ANOVA]. In contrast, combined administration of AP and VU in NSD rats induced a behavioral shift toward an SD-like phenotype, as evidenced by significant deficits PPI that were restricted to the VU + AP condition (**Fig. 7B, D**). PPI deficits were apparent when AP was administered systemically (**Fig. 7B**) [main effect of VU: F(1,28) = 3.81, p = 0.06, $\eta_p^2$ = 0.12; main effect of AP: F(1,28) = 7.91, p = 0.009, $\eta_p^2$ = 0.22; interaction: F(1,28) = 6.26, p = 0.018, $\eta_p^2$ = 0.18; two-way ANOVA]. Similarly, when both AP and VU were infused into the mPFC, robust effects were observed (**Fig. 7D**) [main effect of VU: F(1,31) = 15.01, p = 0.0005, $\eta_p^2$ = 0.33; main effect of AP: F(1,31) = 10.05, p = 0.003, $\eta_p^2$ = 0.24; interaction: F(1,31) = 9.95, p = 0.004, $\eta_p^2$ = 0.24; two-way ANOVA].

These findings demonstrate that while AP is necessary for SD-induced behavioral deficits (as shown by finasteride rescue experiments), it is not sufficient on its own to induce these impairments. Rather, AP requires disrupted chloride homeostasis (elevated $[Cl^-]_i$) to exert its pathological effects.

### SUPPLEMENTARY METHODS

**Animals**. Adult male Sprague Dawley rats (Envigo, Bresso, Italy; 250–320 g at the time of experiments) and adult male and female C57BL/6J mice (Jackson Laboratory, Bar Harbor, ME, USA; 8–12 weeks old, 25–30 g) were used for all experiments. Animals were group-housed (4 animals per cage) in standard polycarbonate cages with corncob bedding and environmental enrichment (nesting material and cardboard tubes). Food (standard rodent chow) and water were provided ad libitum throughout the study. The animal housing room was maintained at 22 ± 0.2°C with 50–60% relative humidity on a 12:12-hour light:dark cycle (lights off at 7:00 PM, lights on at 7:00 AM). Animals were allowed to acclimate to the housing facility for at least 7 days prior to any experimental manipulation. All behavioral, electrophysiological, and biochemical experiments were conducted between 9:00 AM and 5:00 PM (light phase) to minimize circadian variability and ensure consistency with the timing of sleep deprivation procedures, unless otherwise specified. Sample sizes for each experiment were determined a priori through power analyses ($\alpha$ = 0.05, power = 0.80) conducted on preliminary data, with effect sizes estimated from our previous studies (Frau et al., 2008, 2017). Experimenters were blind to treatment conditions during data collection and analysis whenever possible. Every effort was made to minimize the number of animals used and to reduce animal pain, suffering, and distress. All experimental procedures were conducted in accordance with the National Institutes of Health Guide for the Care and Use of Laboratory Animals (8th edition), the European Union directive on the protection of animals used for scientific purposes (EU Directive 2010/63/EU), and Italian legislation (Legislative Decree 26/2014). All protocols were reviewed and approved by the Institutional Animal Care and Use Committee (IACUC) of the University of Cagliari, the Italian Ministry of Health, and the University of Utah (authorization numbers provided upon request).

**Drugs**. The following pharmacological compounds were used in this study:

Allopregnanolone (AP; 3α-hydroxy-5α-pregnan-20-one; Steraloids, Newport, RI, USA) was suspended in 10% (2-hydroxypropyl)-β-cyclodextrin (Sigma-Aldrich, St. Louis, MO, USA) and diluted with sterile 0.9% saline to achieve the desired concentration. For systemic administration, AP was delivered intraperitoneally (IP) at 10 mg/kg. For intracerebral infusions, AP was dissolved in cyclodextrin:Ringer solution (1:5 v/v) to a final concentration of 1 μg/0.5 μl.

Bumetanide (Carbosynth Limited, Berkshire, UK), a selective NKCC1 inhibitor, was suspended in 5% dimethyl sulfoxide (DMSO; Sigma-Aldrich) and diluted with sterile 0.9% saline. BUME was administered systemically (IP) at 25-50 mg/kg, a dose previously shown to inhibit NKCC1 activity in the central nervous system without producing significant diuretic effects (Dzhala et al., 2005).

Finasteride (FIN; Carbosynth Limited), a 5α-reductase inhibitor that blocks the conversion of progesterone to allopregnanolone, was suspended in 10% (2-hydroxypropyl)-β-cyclodextrin and diluted with sterile 0.9% saline. FIN was administered systemically (IP) at 12.5-25 mg/kg or intracerebrally at 0.5 μg/0.5 μl, doses based on previous studies demonstrating effective inhibition of neurosteroid synthesis (Devoto et al., 2012).

VU0463271 (Tocris Bioscience, Bristol, UK), a selective KCC2 inhibitor, was dissolved in 0.5% DMSO and diluted with Ringer solution to a final concentration of 100 μM for intracerebral infusions (0.5 μl/side). This concentration was selected based on published in vitro and in vivo studies demonstrating selective KCC2 inhibition (Delpire et al., 2009).

ANA-12 (Tebubio, Magenta, MI, Italy), a selective TrkB receptor antagonist, was suspended in 5% DMSO and 5% Tween-80 (Sigma-Aldrich) and diluted with sterile 0.9% saline for systemic administration or with Ringer solution for intracerebral infusions. ANA-12 was administered at 1 mg/kg (IP) or 0.5 μg/0.5 μl (intracerebral), doses based on previous studies demonstrating effective TrkB inhibition (Cazorla et al., 2011).

For all systemic administrations, injection volumes were 10 ml/kg for mice and 2 ml/kg for rats. For intracerebral infusions, the volume was 0.5 μl per hemisphere. Vehicle control groups received equivalent volumes of the corresponding vehicle solutions. All drug solutions were freshly prepared on the day of use and protected from light when appropriate.

**Sleep Deprivation (SD)**. using the small-platform-over-water method, a well-validated paradigm that produces near-complete suppression of rapid eye movement (REM) sleep and a substantial reduction in total sleep time in rodents (Cadeddu et al., 2022; Frau et al., 2008, 2017). This method exploits the loss of postural muscle tone during REM sleep, which causes animals to contact the water and awaken, thereby interrupting sleep cycles.

Mice were placed on small circular Plexiglas platforms (4 cm diameter, 6 cm height) inside a tank (50 × 40 × 30 cm) filled with water, leaving 0.5 cm between the water surface and the platform top. Rats were placed on larger platforms (7 cm diameter, 10 cm height) in larger tanks (80 × 60 × 40 cm), with water adjusted to 1 cm below the platform surface. Multiple platforms (4–6 per tank) allowed visual and olfactory contact while preventing physical contact. Water temperature was maintained at 22 ± 1°C. Food and water were provided ad libitum on a wire mesh grid positioned 5 cm above the tank.

Age-matched control animals were housed individually in standard home cages (to match the social isolation experienced by SD animals) in the same experimental room under identical ambient temperature, humidity, and lighting conditions. Control animals had ad libitum access to food and water.

SD lasted 72 h in Sprague Dawley rats, a duration shown to reliably induce prepulse inhibition deficits and manic-like behaviors (Frau et al., 2008, 2017; Gessa et al., 1995), and 24 h in C57BL/6 mice, which display more rapid behavioral responses to SD (Cadeddu et al., 2022). All SD procedures began at 9:00 AM (2 h into the light phase) to standardize circadian timing. Animals were continuously monitored by video throughout SD to ensure procedural compliance and welfare. At the end of SD, animals were immediately transferred to behavioral testing or sacrificed for tissue collection, depending on the experimental endpoint. To control for potential confounding effects of circadian phase, a subset of control animals maintained under an inverted light–dark cycle and was tested at the same clock time (9:00 AM).

**Stereotaxic Surgery and Intracerebral Drug Infusion.** For experiments requiring site-specific drug delivery to the medial prefrontal cortex (mPFC), rats underwent stereotaxic implantation of chronic indwelling guide cannulae. Animals were anesthetized with a combination of fentanyl (0.005 mg/kg, Pfizer, New York, NY, USA) and medetomidine hydrochloride (0.15 mg/kg, Orion Pharma, Espoo, Finland) administered intraperitoneally at a 20:1 ratio. This anesthetic combination provides stable, long-duration anesthesia with minimal respiratory depression and is readily reversible. Depth of anesthesia was confirmed by absence of hindpaw withdrawal reflex and corneal reflex. Body temperature was maintained at 37°C using a feedback-controlled heating pad throughout the surgical procedure.

Rats were placed in a stereotaxic apparatus (David Kopf Instruments, Tujunga, CA, USA) with blunt ear bars to minimize damage to the tympanic membrane. The scalp was shaved, disinfected with povidone-iodine solution, and a midline incision was made to expose the skull. The skull surface was cleaned and leveled by adjusting the incisor bar such that bregma and lambda were in the same horizontal plane. Under aseptic conditions, bilateral craniotomies (approximately 1 mm diameter) were performed using a dental drill. Bilateral stainless steel guide cannulae (22-gauge, 10 mm length; Plastics One, Roanoke, VA, USA) were slowly lowered through the craniotomies and positioned 1 mm dorsal to the target site to minimize tissue damage. Stereotaxic coordinates for the mPFC were: anteroposterior (AP) = +3.0 mm, mediolateral (ML) = ±0.5 mm (angled 10° toward midline), dorsoventral (DV) = −3.0 mm from dura (Paxinos & Watson, 1982). Guide cannulae were secured to the skull using four stainless steel anchor screws (Small Parts Inc., Logansport, IN, USA) and dental acrylic cement (Stoelting Co., Wood Dale, IL, USA). Stainless steel dummy cannulae (Plastics One) were inserted into the guide cannulae to maintain patency and prevent infection. The scalp incision was closed around the implant using surgical sutures.

Following surgery, anesthesia was reversed with atipamezole (1 mg/kg, IP; Orion Pharma), and animals received buprenorphine (0.05 mg/kg, SC) for postoperative analgesia. Animals were placed in clean, warmed recovery cages and monitored continuously until fully ambulatory. Postoperative care included daily monitoring of body weight, food and water intake, and inspection of the surgical site for signs of infection or inflammation. Animals were allowed to recover for 7–10 days before behavioral testing to ensure complete healing and return to baseline body weight.

On test days, drugs were administered via bilateral intracerebral infusions. Dummy cannulae were replaced with 33-gauge internal cannulae (Plastics One) extending 1 mm beyond the guide cannulae and connected to Hamilton syringes (Hamilton Company, Reno, NV, USA) via PE-10 tubing (Intramedic, Becton Dickinson, Franklin Lakes, NJ,

USA). Infusions were delivered at 0.5 µl/min per side using dual microinfusion pumps (CMA 400, CMA Microdialysis, Holliston, MA, USA), with successful delivery confirmed by monitoring air bubble movement. Cannulae remained in place for 2 min post-infusion to allow diffusion and minimize backflow, after which dummy cannulae were reinserted.

Behavioral testing began immediately after infusion for AP or FIN, or 4 h after infusion for ANA-12 to allow maximal TrkB blockade. At the end of experiments, brains were collected, processed, and stained with cresyl violet to verify cannula placements. Only animals with bilateral placements within the prelimbic mPFC (±0.5 mm) were included in analyses, with placement verification performed blind to behavioral outcomes.

**Acute Brain Slice Preparation and Whole-Cell Patch-Clamp Electrophysiology**. Acute cortical brain slices were prepared from adult male Sprague Dawley rats (250–290 g) at 72 hours following the initiation of SD or from age-matched control animals at the equivalent circadian time point. Animals were deeply anesthetized with chloral hydrate (400 mg/kg, IP) and transcardially perfused with ice-cold, oxygenated (95% $O_2$/5% $CO_2$) sucrose-based cutting solution containing (in mM): 75 sucrose, 87 NaCl, 5 KCl, 1.25 $NaH_2PO_4$, 21 $MgCl_2$, 0.5 $CaCl_2$, 25 glucose, and 1.3 ascorbic acid (pH 7.4, osmolarity 300–305 mOsm). The use of sucrose-based cutting solution and elevated magnesium reduces neuronal excitotoxicity and improves slice viability (Ting et al., 2014). Following decapitation, brains were rapidly removed and placed in ice-cold cutting solution. Coronal slices (300 µm thickness) containing the mPFC (prelimbic and infralimbic regions; approximately +3.7 to +2.2 mm from bregma) were prepared using a vibrating microtome (VT1200S; Leica Biosystems, Wetzlar, Germany). Slicing was performed in ice-cold cutting solution continuously bubbled with 95% $O_2$/5% $CO_2$. Slices were immediately transferred to a holding chamber containing artificial cerebrospinal fluid (ACSF) at 32°C for 40 minutes to facilitate recovery, then maintained at room temperature (22–24°C) until recording. ACSF contained (in mM): 126 NaCl, 1.2 KCl, 1.2 $NaH_2PO_4$, 1.2 $MgCl_2$, 2.4 $CaCl_2$, 11 glucose, 18 $NaHCO_3$, and 1.3 ascorbic acid (pH 7.4 when bubbled with 95% $O_2$/5% $CO_2$, osmolarity 304–306 mOsm). Slices were used for recordings within 2–6 hours of preparation.

For recordings, individual slices were transferred to a submerged recording chamber continuously perfused with oxygenated ACSF at 2–3 ml/min at 30–32°C (controlled by an inline heater; Warner Instruments, Hamden, CT, USA). Pyramidal neurons in layer II-III of the prelimbic cortex were visualized using an upright microscope (Axioskop FS 2 plus; Carl Zeiss, Oberkochen, Germany) equipped with infrared differential interference contrast (IR-DIC) optics and a 40× water-immersion objective. Pyramidal neurons were identified based on their characteristic morphology (large pyramidal soma, prominent apical dendrite) and electrophysiological properties (regular spiking pattern, spike frequency adaptation).

Whole-cell patch-clamp recordings were performed using borosilicate glass pipettes (1.5 mm outer diameter, 0.86 mm inner diameter; Sutter Instrument, Novato, CA, USA) pulled on a horizontal puller (P-97; Sutter Instrument) to a resistance of 4–7 MΩ when filled with internal solution. For voltage-clamp recordings of GABA-A inhibitory postsynaptic currents (IPSCs), pipettes were filled with a high-chloride internal solution containing (in mM): 144 KCl, 10 HEPES, 3.45 BAPTA, 1 $CaCl_2$, 2.5 $Mg_2ATP$, 0.25 $Mg_2GTP$ (pH adjusted to 7.2–7.4 with KOH, osmolarity 275–285 mOsm). This high-chloride internal solution was used to enhance the driving force for chloride currents and improve signal-to-noise ratio for IPSC recordings. GABA-A IPSCs were pharmacologically isolated by bath application of the NMDA receptor antagonist D-(-)-2-amino-5-phosphonopentanoic acid (AP5; 100 μM; Tocris Bioscience) and the AMPA receptor antagonist 6-cyano-7-nitroquinoxaline-2,3-dione (CNQX; 10 μM; Tocris Bioscience) to block ionotropic glutamate receptors.

For current-clamp recordings of intrinsic excitability, pipettes were filled with a potassium gluconate-based internal solution containing (in mM): 130 K-gluconate, 10 KCl, 10 HEPES, 0.2 EGTA, 2 $Mg_2ATP$, 0.3 $Na_3GTP$, 10 phosphocreatine-$Na_2$ (pH 7.2–7.4, osmolarity 275–285 mOsm). Current-clamp experiments were performed in standard ACSF without pharmacological blockers.

Recordings were made using an Axopatch 200B amplifier (Molecular Devices, San Jose, CA, USA). Experiments began after the series resistance stabilized (typically within 3–5 minutes of break-in), with acceptable series resistance ranging from 10–35 MΩ. Series resistance and input resistance were continuously monitored throughout experiments using a 5 mV hyperpolarizing voltage step (25 ms duration) applied before each sweep. Cells were excluded from analysis if series resistance changed by >20% during the recording or if the holding current exceeded −200 pA at −70 mV holding potential. Data were filtered at 2 kHz using a four-pole Bessel filter, digitized at 10 kHz using a Digidata 1440A interface (Molecular Devices), and acquired using pClamp 10.2 software (Molecular Devices).

For evoked IPSC recordings, a bipolar stainless steel stimulating electrode (FHC Inc., Bowdoin, ME, USA) was positioned approximately 100 µm rostral to the recording electrode within layer II-III. Stimulation pulses (100–200 µs duration) were delivered at 0.1 Hz using an isolated stimulator (Model 2100; A-M Systems, Sequim, WA, USA). Stimulus intensity was adjusted to evoke IPSCs with amplitudes of 100–300 pA (typically 50–150 µA). Paired-pulse stimulation protocols consisted of two identical stimuli separated by a 50 ms interstimulus interval. Paired-pulse ratios (PPR) were calculated as the amplitude of the second IPSC divided by the amplitude of the first IPSC ($IPSC_2/IPSC_1$), averaged over 20–30 consecutive sweeps during a stable 5-minute baseline period.

For intrinsic excitability measurements, neurons were held at approximately −70 mV (near resting membrane potential) in current-clamp mode. Depolarizing current steps (500 ms duration, 20–400 pA in 20 pA increments, 10 s interstimulus interval) were applied to construct frequency-intensity (F-I) curves. The rheobase (minimum current required to elicit an action potential) was determined using 5 pA increments near threshold. Action potential threshold was defined as the membrane potential at which dV/dt first exceeded 10 mV/ms. Input resistance was calculated from the steady-state voltage response to a −50 pA hyperpolarizing current step. All current-clamp data were corrected for the liquid junction potential (calculated to be approximately −4 mV for the potassium gluconate-based internal solution).

To preserve the native intracellular chloride concentration ($[Cl^-]_i$) and accurately measure the chloride equilibrium potential ($E_{Cl}$) and GABA-A receptor reversal potential ($E_{GABA}$), gramicidin perforated patch-clamp recordings were performed as previously described (Alfonsa et al., 2023; Kyrozis & Reichling, 1995). The gramicidin technique forms pores in the membrane patch that are permeable to monovalent cations ($K^+$, $Na^+$) but not to chloride or other larger anions, thereby providing electrical access to the cell interior while maintaining the endogenous chloride gradient.

Patch pipettes (4–7 MΩ) were filled with a high-chloride internal solution identical to that used for conventional whole-cell recordings (in mM: 144 KCl, 10 HEPES, 3.45 BAPTA, 1 $CaCl_2$, 2.5 $Mg_2ATP$, 0.25 $Mg_2GTP$; pH 7.2–7.4, 275–285 mOsm). Gramicidin D (Sigma-Aldrich) was first dissolved in DMSO to create a stock solution (4 mg/ml), then diluted in the pipette internal solution to a final concentration of 80 µg/ml immediately before use. The gramicidin-containing solution was sonicated for 30 seconds and vortexed thoroughly to ensure complete dispersion. Pipette tips were first front-filled with gramicidin-free internal solution (to facilitate gigaseal formation) by briefly dipping the tip into gramicidin-free solution, then back-filled with gramicidin-containing solution.

Following gigaseal formation (>1 GΩ), gramicidin pores progressively formed in the membrane patch, as evidenced by a gradual decrease in series resistance over 20–40 minutes. Experiments began when series resistance stabilized at approximately 80–120 MΩ, indicating sufficient electrical access. Series resistance was continuously monitored throughout recordings to ensure maintenance of perforated-patch configuration; cells were excluded if series resistance suddenly decreased below 50 MΩ (indicating rupture to whole-cell mode) or if series resistance increased above 150 MΩ (indicating insufficient access).

$E_{GABA}$ measurements were performed in voltage-clamp mode with neurons held at −70 mV. Slow voltage ramps (500 ms duration, ranging from −90 mV to −20 mV, applied every 10 seconds) were delivered at baseline and during bath application of GABA (100 µM; Tocris Bioscience) to activate GABA-A receptors. The GABA-B receptor antagonist CGP35348 (100 µM; Tocris Bioscience) was included in the bath solution to prevent activation of GABA-B receptors. $E_{GABA}$ was determined as the membrane potential at which the difference between the current-voltage relationship in the presence of GABA and the baseline current-voltage relationship (in the absence of GABA) equaled zero. This represents the membrane potential at which there is no net flux of ions through GABA-A receptor channels. $E_{GABA}$ measurements were averaged from 5–10 consecutive ramp protocols during stable GABA application. The Nernst equation was used to calculate the intracellular chloride concentration from the measured $E_{GABA}$, assuming an extracellular chloride concentration of 133 mM (based on ACSF composition).

**In Vivo Two-Photon Chloride Imaging.** Intracellular chloride concentration was measured using the genetically encoded ratiometric biosensor LSSmClopHensor, selectively expressed in cortical pyramidal neurons. Expression of LSSmClopHensor was achieved by viral transfection using a mixture of two viral vectors: AAV9-EF1a-DIO(LSSmClopHensor) and AAV9-CaMKII-Cre. For co-expression, viral mixes were prepared at the following final titers: 8.7 × $10^{11}$ vg/ml for the sensor virus and 2.1 × $10^{11}$ vg/ml for the Cre virus. Adult C57Bl6j mice (4-6 months) were anaesthetized with 2,2,2-tribromoethanol (Avertin) (i.p. 0.02 ml/g body weight), the head of the mouse was gently fixed in a stereotactic frame and the scalp was open above PFC (ML= 0.5 mm and AP= 2 and 3.5 mm anterior to Bregma). A Hamilton NanoFil syringe equipped with a 36G needle (World Precision Instruments) was positioned at the injection sites, and 500 nl of the viral mix (sensor + Cre) was injected at a two depths of 750 and 450 µm below the brain surface at a rate of 100 nl/min. After each injection, the needle was left in place for 1 min to minimize reflux along the injection tract. At the end of the injections, the skin was sutured using sterile surgical thread, and a topical analgesic gel was applied to the incision site. Mice were allowed to recover from anesthesia on a heating pad before returning to their home cage. After surgery, mice were allowed to recover on a heating pad until fully awake and then returned to their home cage. Paracetamol was administered in the drinking water at a dose of 200 mg/kg/day for 2 consecutive days to provide postoperative analgesia and support recovery.

At least three weeks post injections, mice were anesthetized with 2,2,2-tribromoethanol (Avertin) (i.p. 20 µl/g weight) and prepared for acute two-photon imaging. A 4 mm craniotomy was performed over PFC and covered with a 5 mm-coverslip. Imaging was performed immediately after surgery. Mice were positioned underneath a 2-photon microscope. Images were acquired at 5 different excitation wavelengths (800 / 830 / 860 / 910 / 960 nm - some of the 800nm data sets were barely fractionally above noise levels and so were not used for analyses) and collected through green and red emission filters (527/70 nm and 607/70 nm respectively, BrightLine). The imaging was performed at depths ranging from 90 to 300 μm, corresponding to cortical layers 2/3. Post hoc analysis of the images was performed using a graphical user interface (GUI), implemented in MATLAB, that automated finding cell somata, to collect red/green fluorescence ratios at the different wavelengths, and compute intracellular pH and [$Cl^-$] values for each cell, based upon earlier calibration experiments performed using ionophore-permeabilized GL261 cell and neuronal cultures(Sulis Sato et al., 2017). The imaging was performed with a Bruker Ultima microscope, a Coherent Ultra II laser and an Olympus 20X, NA 1.05 objective (water immersion).

**Behavioral Testing.**

*Acoustic Startle Response and PPI.* Sensorimotor gating was assessed using the prepulse inhibition (PPI) of the acoustic startle reflex paradigm, performed as previously described (Bortolato et al., 2004). The startle apparatus (rats: Med Associates, Fairfax, VT, USA; mice: San Diego Instruments, San Diego, CA, USA) consisted of four identical sound-attenuated chambers (56 × 38 × 36 cm), each equipped with a ventilation fan that provided background noise and air circulation. Within each chamber, a transparent Plexiglas cylinder (5 cm inner diameter, 20 cm length for rats; 4 cm diameter, 12 cm length for mice) was mounted on a piezoelectric accelerometric platform that transduced vertical movements of the animal into analog signals. The analog output was digitized and analyzed using a computer-based system. Two separate speakers mounted inside each chamber delivered the background noise and acoustic stimuli, with sound levels calibrated to <1 dB variation across all positions within the cylinder. On the test day, each animal was placed into the Plexiglas cylinder and allowed to acclimate for 5 minutes in the presence of 70 dB broadband white noise, which continued as background throughout the entire session. The test session consisted of three sequential blocks:

- Block 1 (Habituation): Five presentations of the startle pulse alone (120 dB white noise, 40 ms duration) at 15-second intervals to habituate animals to the startle stimulus and reduce initial startle reactivity.

- Block 2 (Test): Fifty trials presented in pseudorandom order with inter-trial intervals ranging from 10–15 seconds (average 12 seconds). Trial types included twelve pulse-alone trials (120 dB, 40 ms); thirty prepulse+pulse trials, consisting of a prepulse (20 ms duration at 74, 78, or 82 dB; 10 trials per intensity) followed 100 ms later by the startle pulse (120 dB, 40 ms); and eight no-stimulus trials (background noise only) to assess baseline movement
- Block 3 (Habituation assessment): Five additional pulse-alone trials identical to Block 1 to assess within-session habituation.

The startle amplitude for each trial was defined as the peak accelerometric response during a 100 ms window beginning at startle stimulus onset. Mean startle amplitude was calculated as the average response to pulse-alone trials in Block 2 (excluding Blocks 1 and 3 to avoid habituation effects). Percent PPI was calculated for each prepulse intensity using the formula:

%PPI = [(mean startle amplitude for pulse-alone trials − mean startle amplitude for prepulse+pulse trials) / mean startle amplitude for pulse-alone trials] × 100

*Novel Object Recognition (NOR).* The test was performed as previously described (Bortolato et al., 2010). The NOR test exploits the natural tendency of rodents to preferentially explore novel objects over familiar ones, providing a measure of declarative memory that depends on intact prefrontal cortex and hippocampal function.

Prior to SD, animals were habituated to the testing environment for two consecutive days. On each habituation day, animals were individually placed in an empty black Plexiglas open-field arena (60 × 60 × 60 cm for rats) and allowed to freely explore for 15 minutes. The arena was cleaned with 70% ethanol and dried between animals to eliminate olfactory cues. Immediately after SD (or at the equivalent time for control animals), each animal was placed in the arena containing two identical objects (e.g., two glass bottles, two plastic cubes) positioned in opposite corners of the arena, equidistant from the center and walls (approximately 15 cm from each wall for rats). Objects were secured to the arena floor with adhesive tape to prevent displacement. Animals were allowed to freely explore both objects for 5 minutes. The time spent actively exploring each object (defined as directing the snout toward the object at a distance ≤2 cm, including sniffing and whisking, but excluding sitting on or near the object without directed attention) was recorded.

Twenty-four hours after the sample phase, animals were returned to the arena. One of the familiar objects from the sample phase was replaced with a novel object that differed in shape, color, texture, and material but was similar in size. The position (left or right corner) of the novel object was counterbalanced across animals. Animals were allowed to explore both objects for 5 minutes, and exploration time for each object was recorded.

A diverse set of objects (constructed from glass, plastic, ceramic, or metal; varied shapes including cylinders, cubes, pyramids; varied colors and surface textures) was used across experiments, with object identities and novel/familiar assignments counterbalanced across treatment groups. All objects were thoroughly cleaned between trials to eliminate olfactory cues. Animals were placed in the arena facing the wall opposite to the objects to prevent initial bias toward one object. All sessions were video-recorded using overhead cameras, and exploration times were scored offline by experimenters blind to treatment conditions using behavioral analysis software (EthoVision XT; Noldus Information Technology, Wageningen, Netherlands).

The recognition index (RI) was calculated as: $RI = [N / (N + F)] \times 100$

where N = time exploring the novel object and F = time exploring the familiar object. An RI significantly greater than 50% indicates preferential exploration of the novel object and thus intact recognition memory.

**Western blotting and immunoprecipitation.** Following completion of behavioral experiments, animals were deeply anesthetized with chloral hydrate (400 mg/kg, IP) and rapidly decapitated. Brains were quickly removed and placed on an ice-cold dissection matrix. The different brain regions (including mPFC, amygdala, nucleus accumbens, and hippocampus) were rapidly dissected using anatomical landmarks, immediately flash-frozen in liquid nitrogen, and stored at −80°C until biochemical processing. All dissections were completed within 3 minutes of decapitation to minimize postmortem protein degradation and dephosphorylation.

For total protein extraction, frozen tissue samples were homogenized on ice using a motorized tissue grinder (Kimble Kontes, Vineland, NJ, USA) in ice-cold HEPES buffer (pH 7.4) containing (in mM): 145 NaCl, 5 KCl, 2 $CaCl_2$, 1 $MgCl_2$, 5 glucose, 5 HEPES, supplemented with protease inhibitor cocktail (cOmplete Mini, EDTA-free; Roche, Basel, Switzerland) and phosphatase inhibitor cocktails 2 and 3 (Sigma-Aldrich) at 1:100 dilution. Homogenates were centrifuged at 12,000 × *g* for 5 minutes at 4°C. The supernatant (total protein fraction) was collected and stored at −80°C.

For membrane-enriched protein fractions, frozen mPFC samples were homogenized in ice-cold 320 mM sucrose/HEPES buffer (pH 7.4) containing 5 mM HEPES, 320 mM sucrose, and protease and phosphatase inhibitors using glass-Teflon homogenizers (Wheaton Science Products, Millville, NJ, USA) with 10 up-and-down strokes at 900 rpm. Homogenates were centrifuged at 900 × *g* for 10 minutes at 4°C to pellet nuclei and debris. The supernatant was collected and centrifuged at 10,000 × *g* for 15 minutes at 4°C to obtain the crude synaptosomal fraction (P2 pellet). The P2 pellet was resuspended in HEPES buffer and centrifuged again at 10,000 × *g* for 15 minutes at 4°C to yield the washed synaptosomal fraction (P2′). The P2′ pellet was resuspended in Cell Lysis Buffer (Cell Signaling Technology, Danvers, MA, USA) containing 20 mM Tris-HCl (pH 7.5), 150 mM NaCl, 1 mM EDTA, 1 mM EGTA, 1% Triton X-100, 2.5 mM sodium pyrophosphate, 1 mM β-glycerophosphate, supplemented with protease and phosphatase inhibitors. Samples were incubated on ice for 30 minutes with periodic vortexing, then centrifuged at 14,000 × *g* for 15 minutes at 4°C. The supernatant (membrane-enriched fraction) was collected and stored at −80°C. Protein concentrations were determined using the DC Protein Assay (Bio-Rad Laboratories, Hercules, CA, USA), a colorimetric assay based on the Lowry method, with bovine serum albumin (BSA) as the standard. Absorbance was measured at 750 nm using a microplate reader (Synergy H1; BioTek Instruments, Winooski, VT, USA).

Phosphorylated KCC2 at threonine 1007 (KCC2-pThr1007) was immunoprecipitated from membrane-enriched fractions using a phospho-specific antibody, as previously described (Pracucci et al., 2023). Rabbit polyclonal anti-KCC2-pThr1007 antibody (custom-generated by the MRC Protein Phosphorylation and Ubiquitylation Unit, University of

Dundee, Scotland, UK; antibody code S961C) was covalently coupled to Dynabeads Protein A (Invitrogen, Carlsbad, CA, USA) according to the manufacturer's protocol. Briefly, 50 µg of antibody was incubated with 1 mg of Dynabeads in phosphate-buffered saline (PBS) with 0.01% Tween-20 for 1 hour at room temperature with rotation. The reaction was quenched with 50 mM Tris-HCl (pH 7.5) for 15 minutes. Antibody-coupled beads were washed extensively and stored at 4°C in PBS with 0.02% sodium azide.

For immunoprecipitation, 250 µg of total protein from membrane-enriched fractions was incubated with 25 µl of antibody-coupled Dynabeads for 3 hours at 4°C with gentle rotation. Beads were washed three times with lysis buffer, and bound proteins were eluted by heating in 4× Laemmli sample buffer (Bio-Rad) containing 5% β-mercaptoethanol at 70°C for 10 minutes. Eluates were analyzed by SDS-PAGE and immunoblotting. To confirm antibody specificity, control experiments included pre-incubation of the antibody with a 10-fold molar excess of the non-phosphorylated KCC2-Thr1007 peptide (University of Dundee), which abolished immunoprecipitation, confirming phospho-specificity.

Protein samples (20–40 µg per lane) were mixed with 4× Laemmli sample buffer containing β-mercaptoethanol, heated at 95°C for 5 minutes, and separated by SDS-polyacrylamide gel electrophoresis (SDS-PAGE) using 4–15% gradient polyacrylamide gels (Criterion TGX Stain-Free Precast Gels; Bio-Rad). Stain-free imaging technology was used to visualize total protein in each lane prior to transfer, providing a loading control independent of housekeeping protein expression (Gürtler et al., 2013). Proteins were transferred to nitrocellulose membranes (0.45 µm pore size; Trans-Blot Turbo Transfer Pack; Bio-Rad) using a semi-dry transfer system (Trans-Blot Turbo; Bio-Rad) according to the manufacturer's protocol (25 V, 1.0 A, 30 minutes).

Following transfer, membranes were blocked in 3% BSA (Sigma-Aldrich) in Tris-buffered saline with 0.1% Tween-20 (TBS-T) for 3 hours at room temperature with gentle shaking. Membranes were then incubated overnight at 4°C with primary antibodies diluted in 3% BSA/TBS-T. Following primary antibody incubation, membranes were washed three times (10 minutes each) in TBS-T and incubated with horseradish peroxidase (HRP)-conjugated secondary antibodies (goat anti-rabbit IgG-HRP or goat anti-mouse IgG-HRP; 1:5000; Jackson ImmunoResearch, West Grove, PA, USA) for 1 hour at room temperature. Membranes were washed three times in TBS-T, and immunoreactive bands were visualized using enhanced chemiluminescence (ECL) substrate (Clarity Western ECL Substrate; Bio-Rad). Chemiluminescent signals were captured using a ChemiDoc XRS+ imaging system (Bio-Rad) with automatic exposure optimization to avoid signal saturation.

Band densities were quantified using Image Lab software version 6.0 (Bio-Rad). For each sample, target protein band densities were normalized to β-actin (loading control) or to total transporter protein (for phospho-transporter measurements) from the same lane. For phosphorylation-specific antibodies, membranes were stripped using Restore Western Blot Stripping Buffer (ThermoFisher Scientific, Waltham, MA, USA) for 15 minutes at room temperature, washed extensively, blocked, and reprobed with antibodies against total (non-phosphorylated) protein to determine phosphorylation stoichiometry.

Representative western blot images are provided in Supplementary Figures S7–11.

The following primary antibodies were used:
Mouse monoclonal anti-β-actin (1:2000; MA5-11869; ThermoFisher Scientific)
Mouse monoclonal anti-β-actin (1:2000; sc-47778; Santa Cruz Biotechnology, Dallas, TX, USA
Rabbit monoclonal anti-KCC2 (1:2000; #94725; Cell Signaling)
Rabbit monoclonal anti-TrkB (1:4000; #4603S; Cell Signaling)
Rabbit polyclonal anti-BDNF (1:6000; ab226843; Abcam)
Rabbit polyclonal anti-GABA-A receptor α1 subunit (1:1000; NB300-191; Novus Biologicals, Centennial, CO, USA)
Rabbit polyclonal anti-GABA-A receptor α4 subunit (1:1000; NB300-193; Novus Biologicals)
Rabbit polyclonal anti-GABA-A receptor δ subunit (1:1000; NB300-200; Novus Biologicals)
Rabbit polyclonal anti-KCC2 phospho-S940 (1:1000; 612-401-E15; Rockland Immunochemicals, Limerick, PA, USA)
Rabbit polyclonal anti-NKCC1 (1:1000; ab303518; Abcam)

Rabbit polyclonal anti-phospho-TrkB (Tyr816) (1:1000; ABN181; Sigma-Aldrich)
Sheep Polyclonal anti-KCC2 phospho-Thr1007 (1:700; S961C; University of Dundee)
Sheep Polyclonal anti-NKCC1 phospho-Thr203/Thr207/Thr212 antibody (1:700; S763B; University of Dundee).

**Immunofluorescence.** Following SD exposure, mice were deeply anesthetized with isoflurane and transcardially perfused with PBS. Brains were harvested, post-fixed in 4% paraformaldehyde (PFA) for 48 h, cryoprotected in 30% sucrose, and sectioned coronally at 20 μm thickness. Brain sections were incubated in TBS-A (TBS containing 0.1% Triton X-100), followed by blocking in TBS-B (TBS-A supplemented with 2% BSA). Sections were then incubated overnight at 4°C with a polyclonal anti-BDNF antibody (NB100-98682, Novus Biologicals) diluted 1:200 in TBS-B. On the following day, sections were washed twice in TBS-A and once in TBS-B before incubation with the appropriate secondary antibody diluted in TBS-B for 1 h at room temperature in the dark (Goat anti-rabbit IgG Alexa Fluor 647; ThermoFisher Scientific). Nuclear counterstaining was performed by adding DAPI (Sigma-Aldrich) to the secondary antibody solution. Sections were subsequently washed in TBS and mounted using Fluoroshield mounting medium (Sigma-Aldrich). Images were acquired with a Nikon C1 confocal microscope.

**Statistical Analyses**. Prior to statistical testing, data were assessed for normality using the Kolmogorov–Smirnov test and for homogeneity of variance using Bartlett's test. Data that violated assumptions of normality were log-transformed or, when appropriate, analyzed using non-parametric tests. Unless otherwise specified, all measurements were obtained from independent biological samples, with each animal contributing a single data point per outcome. Repeated measurements were used only in experimental designs that explicitly included a within-subject factor (e.g., electrophysiological input–output curves).

Comparisons between two independent groups were performed using unpaired two-tailed Student's *t*-tests. Comparisons involving more than two groups or multiple factors were analyzed using multi-way analysis of variance (ANOVA), with main effects and interactions considered significant at $p < 0.05$. When significant effects were detected, post hoc pairwise comparisons were conducted using Tukey's test, with Spjøtvoll–Stoline correction applied when group sizes were unequal. For repeated-measures designs, sphericity was assessed using Mauchly's test, and Greenhouse–Geisser corrections were applied when violations were detected.

Effect sizes were calculated for all primary statistical analyses using JASP (v. 0.19.3) and are reported alongside inferential statistics. Cohen's *d* was reported for t tests, and partial eta squared ($\eta_p^2$) was reported for ANOVA-based analyses.

Prepulse inhibition (PPI) data were initially analyzed using three-way ANOVA with sleep condition, drug treatment, and prepulse intensity as factors. As no significant interactions involving prepulse intensity were observed, PPI values were averaged across prepulse levels for each subject, yielding a single mean %PPI per animal. Subsequent analyses were therefore performed using two-way ANOVA (sleep condition × drug treatment) on independent subject means.

For all analyses, the threshold for statistical significance was set at $\alpha = 0.05$ (two-tailed). Exact p-values and corresponding effect sizes are reported in the Supplementary Results. All data collection, behavioral scoring, and quantitative analyses were performed by experimenters blinded to treatment conditions and experimental hypotheses.
